# Discovery of a celecoxib binding site on PTGES with a cleavable chelation-assisted biotin probe

**DOI:** 10.1101/782532

**Authors:** David K. Miyamoto, Hope A. Flaxman, Hung-Yi Wu, Jinxu Gao, Christina M. Woo

## Abstract

The coxibs are a subset of non-steroidal anti-inflammatory drugs (NSAIDs) that primarily target cyclooxygenase-2 (COX-2) to inhibit prostaglandin signaling and reduce inflammation. However, mechanisms to inhibit other members of the prostaglandin signaling pathway may improve selectivity and reduce off-target toxicity. Here, we report a novel binding site for celecoxib on prostaglandin E synthase (PTGES), an enzyme downstream of COX-2 in the prostaglandin signaling pathway, using a cleavable chelation-assisted biotin probe **6**. Evaluation of the multi-functional probe **6** revealed significantly improved tagging efficiencies attributable to the embedded picolyl functional group. Application of the probe **6** within the small molecule interactome mapping by photo-affinity labeling (SIM-PAL) platform using photo-celecoxib as a reporter for celecoxib identified PTGES and other membrane proteins in the top eight enriched proteins from A549 cells. Carbonic anhydrase 12, a known protein target of celecoxib, was also enriched. Four binding sites to photo-celecoxib were additionally mapped by the probe **6**, including a binding site with PTGES. The binding interaction with PTGES was validated by competitive displacement with celecoxib and known PTGES inhibitor licofelone. The binding site of photo-celecoxib on PTGES enabled the development of a structural model of the interaction and will inform the design of new selective inhibitors of the prostaglandin signaling pathway.

## Main text

Inflammation is a major immune response to injury or infection that leads to short-term symptoms of swelling and pain and, when dysregulated, long-term diseases including arthritis, autoimmune disorders, and neurodegeneration. Induction of the acute inflammatory response is primarily mediated by prostaglandin signaling.^1^ The prostaglandins are produced by the cyclooxygenases COX-1 and COX-2, which transform arachidonic acid into prostaglandin H_2_ (PGH_2_) for further tailoring by prostaglandin synthases (Figure 1A). The specific inhibition of prostaglandin E_2_ (PGE_2_) production and signaling has been an anti-inflammatory target since the development of the first non-steroidal anti-inflammatory drug (NSAID), aspirin, and the subsequent introduction of selective COX-2 inhibitors known as the “coxibs”.^2^ While therapeutic inhibition of COX-1 and COX-2 is associated with gastrointestinal toxicity, the selective COX-2 inhibitor rofecoxib was withdrawn due to cardiovascular toxicity.^3^ This cardiovascular toxicity may arise from the simultaneous suppression of PGE_2_ and the cardiovascular protectant PGI_2_ by selective COX-2 inhibitors.^4^ As a result, identification of PGE_2_-selective inhibitors by targeting the inducible prostaglandin E synthase (PTGES) has been the focus of several efforts.^5–8^ Here, we report the discovery of a direct interaction between PTGES and celecoxib using binding site hotspot mapping with an isotopically-coded cleavable chelation-assisted biotin probe (CBPA, **6**). Chemical proteomics methods have enabled the unbiased identification of protein targets, altered pathways, and recently non-covalent binding site hotspots of small molecules in the cellular proteome.^9–11^ Recently, we reported a platform termed small molecule interactome mapping by photo-affinity labeling (SIM-PAL) that combines cellular photo-affinity labeling (PAL) and chemical enrichment methods with isotope-targeted mass spectrometry (MS) to map the binding sites of several NSAIDs, including celecoxib (**1**).^11^ Derivatization of celecoxib (**1**) with a diazirine-based PAL group guided by the structure–activity relationship of COX-2 binding yielded photo-celecoxib (**2**) as a reporter for celecoxib (**1**, Figure 1B).^12^ Application of photo-celecoxib (**2**) to live cell labeling, followed by photo-conjugation and copper-catalyzed azide-alkyne cycloaddition (CuAAC) with the multifunctional cleavable biotin azide probe (CBA, **5**) enabled the selective enrichment and isotopic recoding of celecoxib-conjugated peptides for analysis by isotope-targeted MS (Figure 1C). Targeted database searching yielded binding site maps of photo-celecoxib (**2**) to COX-2 in vitro and in the global proteome in Jurkat and K562 cells.^11^

**Figure 1.**
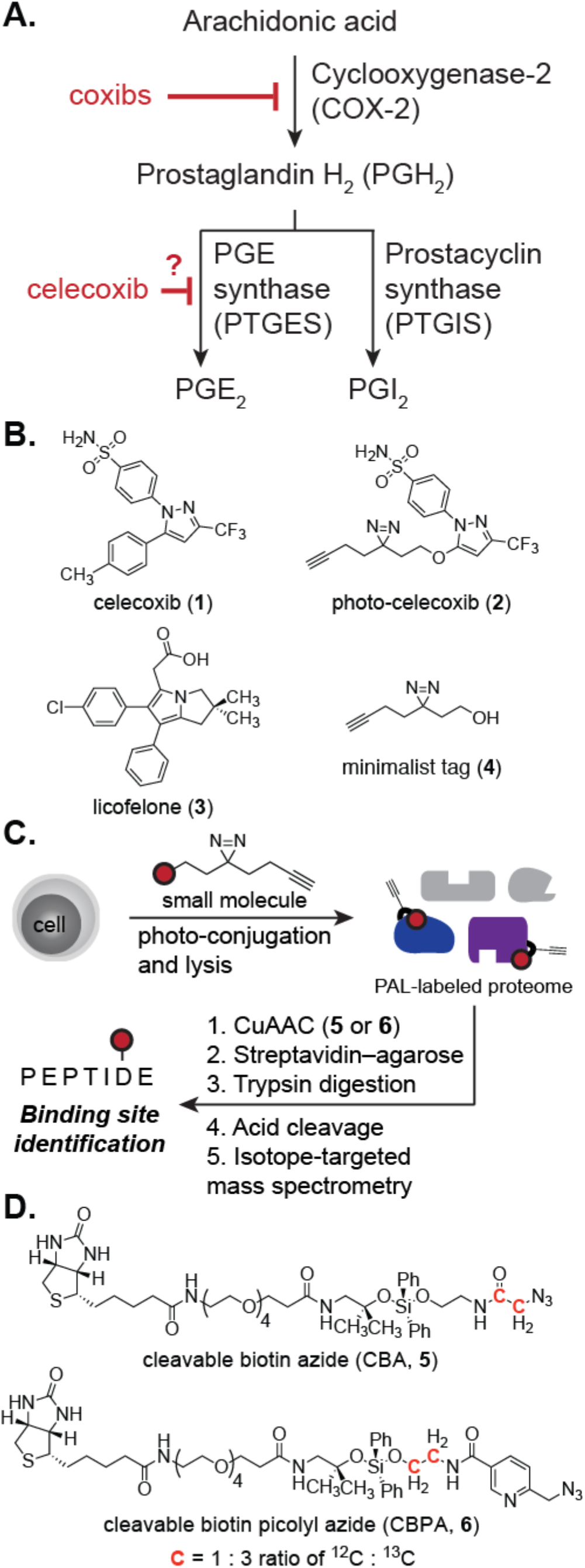
Overview of the structural characterization of celecoxib binding interactions by binding site hotspot mapping. **A.** The prostaglandin signaling pathway is inhibited by the coxibs. **B.** Structures of celecoxib (**1**), photo-celecoxib (**2**), licofelone (**3**), and the minimalist tag (**4**). **C.** Workflow for SIM-PAL, resulting in binding site identification. **D.** Structures of the cleavable biotin azide probe **5** and the cleavable biotin picolyl azide **6**.

In the course of these studies, we found that CuAAC with a standard biotin azide often yielded higher overall anti-biotin signal than CBA **5**, which may have been caused by incomplete CuAAC with the small molecule-labeled proteome. Inspired by chelation-assisted catalysis,^13^ we thus designed CBPA **6** to improve the throughput of binding site hotspot mapping (Figure 1D, Scheme S1). CBPA **6** incorporates a picolyl group adjacent to the azide to promote copper-chelation and thus increase CuAAC efficiency in addition to a biotin enrichment handle and diphenylsilane linker. The silane linker is cleaved under mild acid hydrolysis (2% formic acid, 24 °C),^14–16^ conditions that are highly compatible with downstream MS analysis.

We initially evaluated CBPA **6** against a HEK293T cell lysate labeled with homo-propargyl glycine (HPG, 1 mM). The probe, copper sulfate, and ligand concentrations were optimized for CuAAC with CBPA **6**, and with the optimized probe–ligand ratio, a direct comparison of the CuAAC efficiency with biotin-(PEG)_3_-azide and the cleavable biotin probes **5** and **6** was performed (Figure 2A). HPG-labeled HEK293T lysates were treated with a pre-mixed solution of the CuAAC reagents to a final concentration of 100 µM of the biotin probe, 250 µM CuSO_4_, 250 µM THPTA, and 2.5 mM sodium ascorbate for 1.5 h and the degree of biotin tagging was visualized by streptavidin–IR680LT. CBPA **6** showed a 7-fold increase in CuAAC efficiency compared to CBA **5**, and reached 70% of the tagging efficiency of the standard biotin-(PEG)_3_-azide (Figure 2A, 2B, S1).

**Figure 2.**
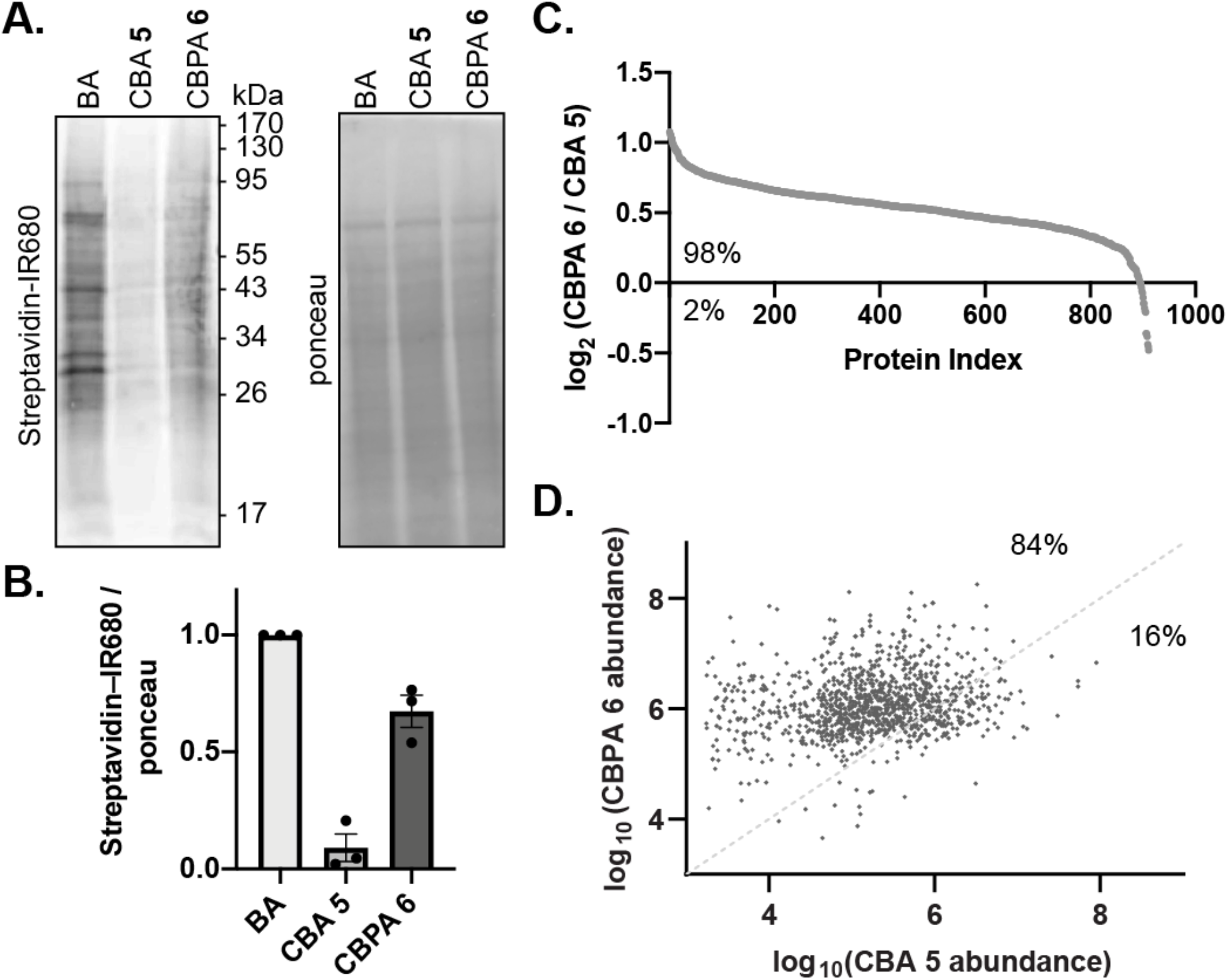
Evaluation of biotin probes against an HPG-labeled HEK293T lysate. **A.** Western blot of HPG-labeled lysates after click chemistry with a biotin-azide probe. BA = biotin-(PEG)_3_-azide, CBA = cleavable biotin azide, CBPA = cleavable biotin picolyl azide. **B.** Relative quantification of the streptavidin–IR680 signal normalized to total protein input (ponceau) across three biological replicates. **C.** Comparison of CBPA **6** or CBA **5** by TMT-based quantitative proteomics following CuAAC, enrichment, and trypsin digestion of HPG-labeled lysates. **D.** Label-free quantification of unique HPG-labeled peptides tagged by CBA **5** or CBPA **6** following acid cleavage and recovery of the modified peptide from the streptavidin–agarose resin. The percentages given indicate the fraction of HPG-labeled peptides lying above and below the line of unity.

The observed increase in CuAAC efficiency by CBPA **6** was likewise observed by quantitative proteomics. From a total of 911 proteins enriched by both biotin probes **5** and **6** across two biological replicates, 98% of the proteins were more enriched when treated with CBPA **6** (Figure 2C, Table S2). Importantly, an improvement in site identification was additionally observed. Spectral counting of isotopically-coded HPG-embedded peptides showed 7,524 PSMs tagged with CBPA **6** as compared to 3,441 PSMs tagged by CBA **5**, representing a greater than 50% increase in PSMs following enrichment by CBPA **6** (Table S3). Label-free quantification was subsequently evaluated by integration of the abundance values for unique HPG-containing peptides detected across both samples and showed that 84% of these peptides were enriched to a greater degree by CBPA **6** (Figure 2D). We additionally compared the enrichment efficiency using CuAAC conditions that were individually optimized to CBA **5** and CBPA **6** (Table S1). Using CuAAC conditions optimized for each probe, we found that the degree of biotin tagging still favored CBPA **6** and, although protein quantification values were similar, 87% of HPG-labeled peptides were more enriched by CBPA **6** than by CBA **5** (Figure S2).

Encouraged by the improvement in enrichment yields using CBPA **6**, we next investigated the application of CBPA **6** to binding site maps produced by photo-celecoxib (**2**). The human epithelial lung carcinoma cell line A549 is a frequently used model for prostaglandin signaling due to the ability to produce a variety of prostaglandins under stimulation.^17^ Photo-celecoxib (**2**) and celecoxib (**1**) display similar inhibitory properties for COX-2^11^ and both exert anti-proliferative effects in A549 cells at micromolar concentrations (Figure S3). A549 cells were therefore treated with 10 µM photo-celecoxib (**2**) for 2 h followed by photo-irradiation to capture the binding interactions. The treated cells were collected, lysed in 1% RapiGest in PBS, and tagged with CBPA **6**. The biotinylated proteins were enriched on streptavidin–agarose beads, followed by on-bead tryptic digestion for TMT labeling and quantitative proteomics. Acid cleavage of CBPA **6** released the photo-celecoxib-conjugated bead-bound peptides for direct analysis of the binding sites by LC-MS/MS.

Quantitative proteomics analysis of the peptides released by on-bead trypsin digestion revealed 666 enriched proteins (3 or more unique peptides, Figure 3A, Table S4). We evaluated potential targets based on comparison to the “diazirine photoome” in A549 cells and found eight proteins in the photo-celecoxib dataset enriched greater than any protein from the “diazirine photoome”.^18^ Alternatively, potential targets could be identified by competitive displacement of the photo-celecoxib proteome with the parent molecule. These eight proteins are intriguingly all membrane proteins localized to the endoplasmic reticulum or mitochondrial membrane and are involved in translocation of proteins across these membranes (e.g., KDELR1, TRAM1, TMEM97, TIMM17B, TMEM245) or redox homeostasis (e.g., MGST1, PTGES, CYB5B, Figure 3B). Notably, cytochrome b5 type B (CYB5B) was enriched along with an additional 15 subunits of several cytochrome complexes (highlighted in blue, Figure 3A). Cytochrome P450 2S1 (CYP2S1), which may be involved in celecoxib metabolism,^19^ was also identified with two unique peptides. The identification of proteins that are part of complexes is in line with previous observations using PAL-based enrichment and may represent conformational flexibility that is captured by PAL.^11, 20^ The cytochrome complex was enriched along with a known target carbonic anhydrase 12 (CA12, highlighted in green, Figure 3A).^21^ Of the significantly enriched proteins, we were particularly interested in the identification of PTGES due to its role downstream of COX-2 in the prostaglandin biosynthetic pathway (highlighted in red, Figure 3A). However, COX-2 itself was not observed in the quantitative proteomics data potentially due to the need to induce COX-2 expression in A549 cells with phorbol 12-myristate 13-acetate (PMA) stimulation.^17^ Indeed, COX-2 was only observed in A549 cells after stimulation with PMA where it was susceptible to enrichment by photo-celecoxib (**2**) and competitively displaced by celecoxib (**1**, Figure S4).

**Figure 3.**
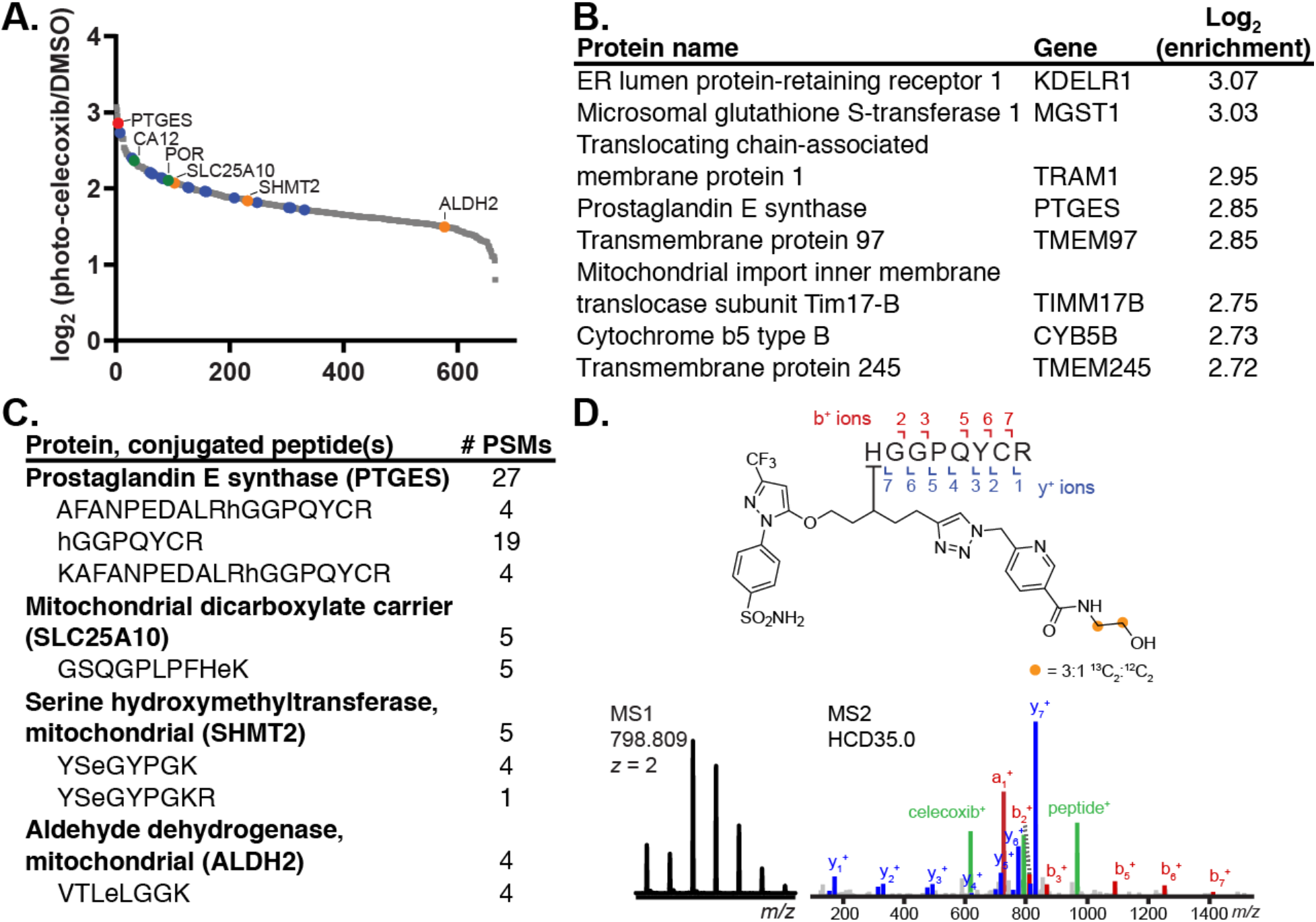
Binding site hotspot mapping of photo-celecoxib (**2**) in A549 cells. **A.** Quantitative proteomics data of photo-celecoxib (**2**) normalized to DMSO control. Proteins highlighted in red = enriched protein and binding site (PTGES); orange = binding site identified; green = previously identified target of celecoxib; blue = members of the cytochrome complex. **B.** Top eight proteins enriched from A549 cells by photo-celecoxib (**2**). **C.** Peptides observed directly conjugated to photo-celecoxib (**2**) and the frequency of peptide spectral matches (PSMs). Assigned conjugation site is represented by a lower-case letter. **D.** Example PSM assignment of a binding site from PTGES.

We next evaluated the binding sites observed upon acid cleavage of CBPA **6**, recovery of the photo-celecoxib-conjugated peptide, and filtration of the assignments through the IsoStamp software.^22^ The isotopic code incorporated into CBPA **6** facilitated detection of photo-celecoxib-labeled peptides. Binding sites were identified on four proteins following manual validation (Figure 3C; highlighted in red/yellow, Figure 3A, Table S5). These binding sites corresponded to proteins that were significantly enriched (PTGES) and enriched above background (SLC25A10, SHMT2, ALDH2). ALDH2 may be involved in the metabolism of celecoxib.^19^ Spectral counting of each binding site is in rough correlation with the degree of protein enrichment, and may represent a mechanism to approximate the degree of protein enrichment from binding site data alone. However, these data illustrate that the identification of a binding site to a target protein is not inherently restricted to highly enriched proteins and is dependent on many other factors, including the overall abundance of the target protein, the amino acid preferences of the PAL chemistry, and the existence of the binding site within a protease-accessible region of the protein. Evaluation of a broader array of proteases and PAL chemistries across cell lines will extend the ability to access a wider range of binding sites by MS.

The binding site on PTGES was discovered with a total of 27 photo-celecoxib-conjugated PSMs across two biological replicates. This binding site was reproducibly detected across four biological replicates enriched by CBPA **6**. An example peptide spectral match (PSM) of photo-celecoxib (**2**) conjugated to PTGES His-53 is shown in Figure 3D. PTGES is a membrane-associated synthase that performs the terminal biosynthetic processing of PGH_2_ to PGE_2_.^23^ As PTGES exists as a trimeric complex in the microsomal membrane,^8^ the occurrence of multiple binding events may contribute to the frequent binding site observation. The expression and activity of PTGES is inducible by pro-inflammatory stimuli and is functionally coupled to COX-2.^24^ Although celecoxib (**1**) exhibits limited inhibitory activity against PTGES relative to COX-2,^25^ the cellular treatments performed here are on a similar order of magnitude as the C_max_ for celecoxib in humans (C_max_ = 1.8 µM).^26^ We therefore evaluated the binding interaction between PTGES and celecoxib by competitive displacement. The enrichment of PTGES in the presence of 10 µM photo-celecoxib (**2**) was validated by Western blot and observed to be competed by the parent compound (10x concentration, Figure 4A). Furthermore, several small molecule inhibitors have been developed against PTGES, including licofelone (**3**).^27^ Licofelone (**3**) likewise competitively displaced 10 µM photo-celecoxib (**2**) from PTGES (Figure 4A) at a 10x concentration.

**Figure 4.**
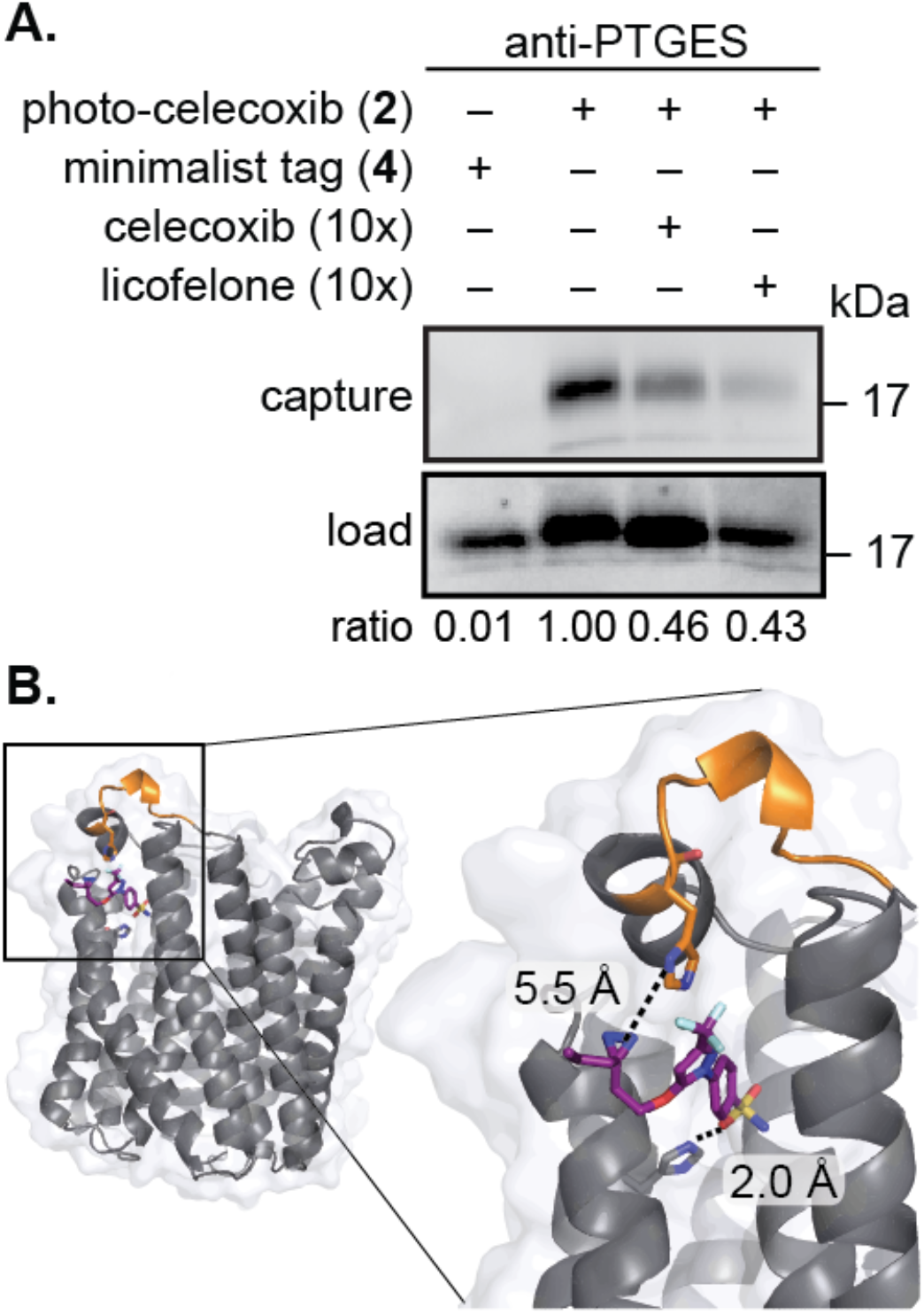
Validation and docked model of PTGES with photo-celecoxib (**2**) in A549 cells. **A.** Western blot for PTGES before and after competitive displacement of photo-celecoxib (**2**) by celecoxib (**1**) or licofelone (**3**) following photo-irradiation and enrichment from A549 cells. **B.** Docked model of PTGES and photo-celecoxib (**2**) from PDB 4YK5. The binding site HGGPYR is highlighted in orange. The zoomed image shows the distance between His-53 and the diazirine on photo-celecoxib (**2**) in the docked structure.

With the binding site in hand, we sought to develop a model of photo-celecoxib (**2**) bound to PTGES. Structures of inhibitors bound to PTGES display two distinct binding modes, one of which is in close proximity to the observed binding site.^6–8^ Photo-celecoxib (**2**) was modeled into the structure of PTGES (PDB 4YK5)^8^ by simulated annealing with distance minimization using Amber. Based on prior measurements of the diazirine labeling radius,^20^ a 9 Å distance restraint between the diazirine and His-53 on PTGES was used to determine the feasibility of labeling in the resulting structures. Following simulated annealing, the lowest energy structure was minimized to yield the model displayed in Figure 4B. In the model, photo-celecoxib (**2**) occupies a binding site between two monomers of PTGES that appears to be stabilized by hydrogen bonding between His-113 and the sulfonamide on photo-celecoxib (**2**, Figure 4B).

In conclusion, we designed and synthesized an isotopically-coded acid-cleavable chelation-assisted probe **6** to improve enrichment efficiencies for binding site mapping. Optimization of the CuAAC protocol with CBPA **6** improved protein enrichment for 98% of proteins and modification site identification for 84% of peptides compared to CBA **5**, using HPG-labeled cell lysates. Enrichment using CBPA **6** with photo-celecoxib (**2**) revealed a pool of 666 proteins of which the top eight enriched proteins were membrane-bound, including PTGES, a protein downstream of COX-2 in the prostaglandin signaling pathway. Binding site hotspot mapping further revealed four unique binding sites, including a binding site on PTGES with photo-celecoxib (**2**) conjugated to His-53. The observed binding event was confirmed by competitive displacement by celecoxib (**1**) and known PTGES inhibitor licofelone (**3**). The interaction between photo-celecoxib and PTGES was modeled using Amber. These insights will assist in the development of new inhibitors of the prostaglandin signaling pathway and provide an improved biotin probe for the versatile discovery of the protein and binding site interactions of bioactive small molecules.

## Associated Content

Supporting figures and procedures (PDF)

Supplementary tables 2–5 (Excel)

The mass spectrometry proteomics data have been deposited to the ProteomeXchange Consortium via the PRIDE partner repository with the dataset identifier PXD014299.

## Acknowledgements

We thank B. Norman and M. Labenski for insightful comments and B.Budnik (Harvard Proteomics Resource Laboratory) for acquiring MS data. Financial support from the Burroughs Wellcome Fund (C.M.W.), the Ono Pharma Foundation (C.M.W.), the Sloan Foundation (C.M.W.), the National Science Foundation GRFP (H.A.F.), and Harvard University is gratefully acknowledged.

## Supporting Information Available

This material is available free of charge *via* the Internet.

