## Supporting Information for "Discovery of a celecoxib binding site on PTGES with a cleavable chelation-assisted biotin probe"

*ACS Chemical Biology*

Supporting Information

### Index

**Additional Experimental Information** ........ **S2**

**Scheme S1.** Synthesis of the biotin picolyl azide probe **6**

**Table S1.** Optimized CuAAC conditions for biotin azide probes

**Figure S1.** Comparison of biotin–azide, CBA **5**, and CBPA **6** CuAAC efficiency by streptavidin-HRP Western blot
**Figure S2.** Evaluation of CBA and CBPA with individually optimized CuAAC conditions
**Figure S3.** Cell viability assay (MTT) for celecoxib (**1**) and photo-celecoxib (**2**) in A549 cells
**Figure S4.** Interaction of photo-celecoxib with COX-2 in A549 cells

**General Experimental Procedures** ........ **S8**
**Chemical Materials** ........ **S8**
**Biological Materials** ........ **S8**
**Cell Culture Materials** ........ **S8**
**Chemical Instrumentation** ........ **S8**
**Experimental Procedures with Cell Lysates** ........ **S10**
**Experimental Procedures with Whole Cells** ........ **S12**
**Mass Spectrometry Procedures** ........ **S13**
**Data Analysis Procedures** ........ **S13**
**Structural Modeling Procedures** ........ **S14**
**Synthetic Procedures** ........ **S15**
**Catalog of Nuclear Magnetic Resonance and Infrared Spectra** ........ **S20**
**Catalog of Unique Binding Site Peptide Spectral Matches** ........ **S32**
**Bibliography** ........ **S39**

#### Additional Experimental Information.

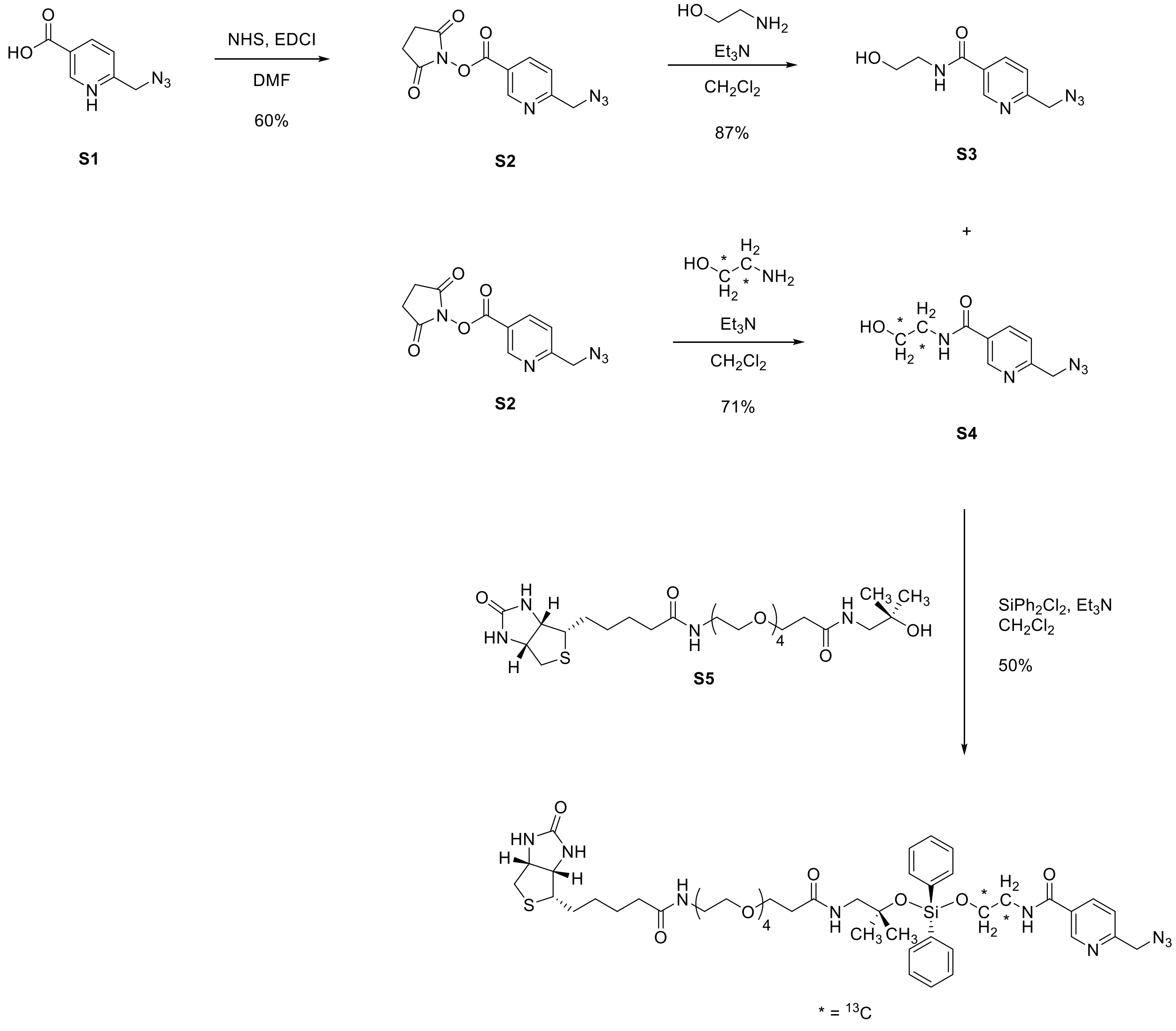


cleavable biotin picolyl azide probe **6** ($^{13}$C, 75%)

**Scheme S1.** Synthetic scheme for the cleavable biotin picolyl azide probe **6**.

| | Final concentration | |
|---|---|---|
| Reagent/Time | **CBA 5** | **CBPA 6** |
| reaction time | 3 hours | 1.5 hours |
| probe | 200 µM | 100 µM |
| $CuSO_4$ | 300 µM | 250 µM |
| THPTA | 600 µM | 250 µM |
| sodium ascorbate | 2.5 mM | 2.5 mM |

**Table S1.** Optimized CuAAC conditions for the cleavable biotin azide probe **5** and the cleavable biotin picolyl azide probe **6**.

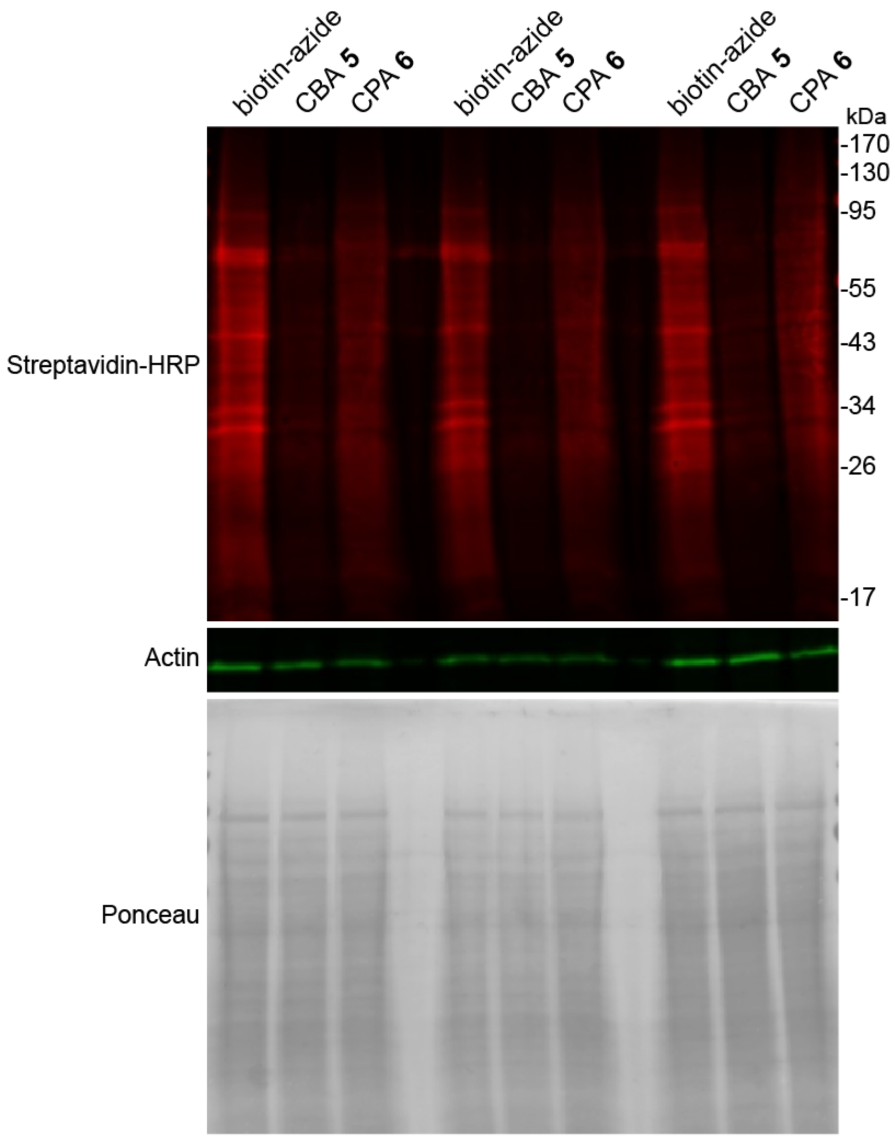


**Figure S1.** Comparison of biotin–azide, CBA **5**, and CBPA **6** CuAAC efficiency by streptavidin-HRP Western blot. CuAAC was performed on HPG-labeled lysates using conditions optimized for CBPA **6** (Table S1).

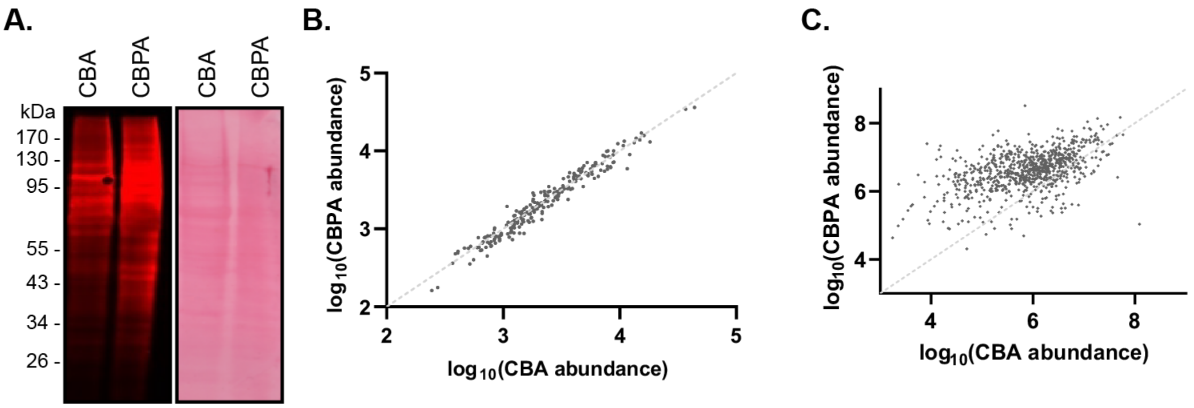


**Figure S2.** Evaluation of the biotin probes CBA **5** and CBPA **6** using individually optimized conditions for each probe as shown in Table S1. **A.** Streptavidin–HRP evaluation of CBA **5** and CBPA **6** using individually optimized conditions. **B.** Quantitative proteomics of HPG-labeled lysates after enrichment showing similar enrichment of CBA **5** and CBPA **6**. **C.** Label-free quantification of HPG-containing peptides enriched by both CBA **5** and CBPA **6**.

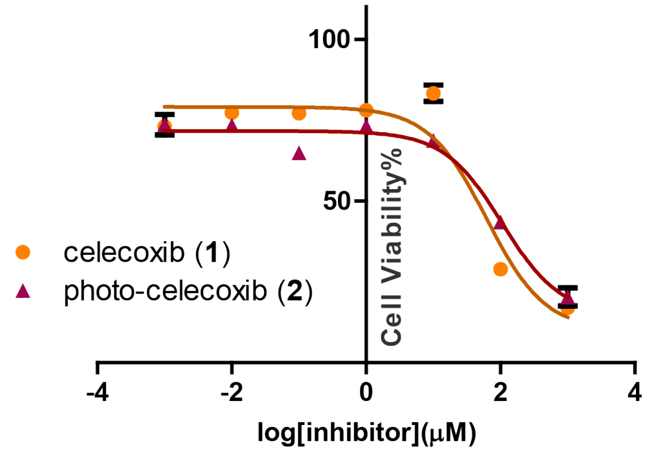


| Inhibitor | $IC_{50}$ (μM) |
|---|---|
| celecoxib (**1**) | 44.65 |
| photo-celecoxib (**2**) | 64.55 |

**Figure S3.** Cell viability assay (MTT) for celecoxib (**1**) and photo-celecoxib (**2**) in A549 cells.

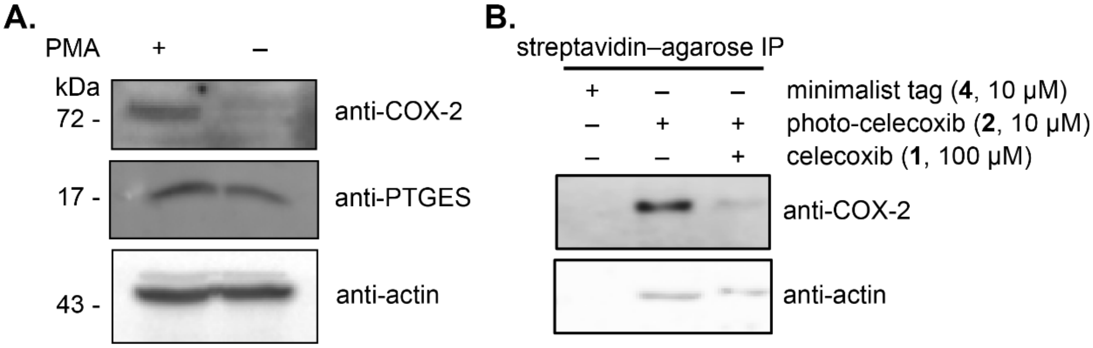


**Figure S4.** Interaction of photo-celecoxib with COX-2 in A549 cells. **A.** Western blot for COX-2, PTGES, and actin with and without stimulation with PMA in A549 cells. A549 cells were stimulated with PMA (10 nM) for 6 h at 37 ℃. Cells were lysed and protein expression was analyzed by Western blot. **B.** Streptavidin–agarose enrichment of labeled proteins from stimulated A549 cells. Cells were treated with minimalist tag (**4**), or photo-celecoxib (**2**), or photo-celecoxib (**2**) and competed with a 10x concentration of celecoxib (**1**) for 2 h at 37 ℃. Cells were then irradiated and tagged with the cleavable biotin azide **5** and biotinylated proteins were enriched on streptavidin–agarose resin. The resin (capture) was analyzed by Western blot.

**General Experimental Procedures.** All reactions were performed in single-neck, flame-dried, round-bottomed flasks fitted with rubber septa under a positive pressure of argon, unless otherwise noted. Air- and moisture-sensitive liquids were transferred via syringe or stainless steel cannula. Organic solutions were concentrated by rotary evaporation at 30–33 °C. Normal and reverse phase flash-column chromatography was performed as described by Still and co-workers.[1] Normal phase purifications employ silica gel (60 Å, 40–63 μm particle size) purchased from Silicycle (Quebec, Canada). Analytical thin-layer chromatography (TLC) was performed using glass plates pre-coated with silica gel (0.25 mm, 60 Å pore size) impregnated with a fluorescent indicator (254 nm). TLC plates were visualized by exposure to ultraviolet light (UV), iodine ($I_2$), and/or submersion in *p*-anisaldehyde followed by brief heating with a heat gun (10–15 s).

**Chemical Materials.** Commercial solvents and reagents were used as received with the following exceptions. Dichloromethane and *N,N*-dimethylformamide were purified according to the method of Pangborn and co-workers.[2] Triethylamine and ethanolamine were distilled from calcium hydride under an atmosphere of nitrogen immediately before use. RapiGest was prepared according to the method of Lee and co-workers.[3] 6-(Azidomethyl)nicotinic acid **S1** was synthesized according to the method of Ting and co-workers.[4] Biotin-CA($PEG)_4$-alcohol **S5** was synthesized according to the method of Tirrell and co-workers.[5] Photo-celecoxib (**2**) was synthesized according to the method of Woo and co-workers.[6] Minimalist tag (**4**) was synthesized according to the method of Yao and co-workers.[7]

**Biological Materials.** PTGES polyclonal antibody (Thermo Fisher Scientific, #PA5-51036) and COX-2 polyclonal antibody (Cell Signaling Technology, #4842) were diluted to 1:1,000 in 3% BSA/TBST for Western blot detection. β-actin monoclonal antibody (Santa Cruz Biotechnology, #SC-47778), high sensitivity streptavidin-HRP (Thermo Fisher Scientific, #21130), and IRDye 680LT Streptavidin (LI-COR Biosciences, #926-68031) were diluted to 1:10,000 in 3% BSA/TBST for Western blot detection. Protease inhibitor tablets (Roche EDTA-free cOmplete tablets, Sigma-Aldrich, # 11836170001) were resuspended as a 25x stock in 2 mL PBS and stored at −20 ºC. L-homopropargylglycine (Click Chemistry Tools, #1067) was resuspended as a 100 mM stock in water and stored at 4 ºC. Phorbol 12-myristate 13-acetate (Sigma Aldrich, #P1585) was resuspended as a 10 μM stock in DMSO and stored at –20 ºC. Biotin-($PEG)_3$-azide (Sigma Aldrich, #762024) was resuspended as a 10 mM stock in DMSO and stored at –20 ºC. Tris(3-hydroxypropyltriazolylmethyl)amine (THPTA, Sigma Aldrich, #762342) was resuspended as a 10 mM stock in DMSO and stored at –20 ºC. Licofelone was purchased from Santa Cruz Biotechnology (#SC-207826) and resuspended as a 50 mM stock in DMSO and stored at –20 ºC. For LC-MS/MS analysis, proteins were digested with sequencing grade trypsin (Promega, # V5111). M-PER was obtained from Thermo Scientific (#78501). Mini Bio-Spin chromatography columns were obtained from Bio-Rad (#732-6207). Streptavidin–agarose beads were obtained from Thermo Scientific (#20353) and washed with PBS prior to use. BCA reagent A (G-Biosciences, #786-846) and BCA reagent B (G-Biosciences, #786-848) were mixed in a 50:1 ratio and used to measure protein concentrations of cell lysates.

**Cell Culture Materials.** HEK 293T and A549 cell lines were obtained from the American Type Culture Collection (ATCC) and maintained in DMEM medium supplemented with 10% fetal bovine serum (FBS) and 1% penicillin/streptomycin at 37 ˚C and 5% $CO_2$ in a water-saturated incubator. HPG-incorporated HEK 293T cells were grown in leucine- and methionine-free medium (Thermo Scientific, #30030).

**Chemical Instrumentation.** Proton nuclear magnetic resonance spectra ($^1H$ NMR) were recorded at 400 or 500 MHz at 24 °C, unless otherwise noted. Chemical shifts are expressed in parts per million (ppm, δ scale) downfield from tetramethylsilane and are referenced to residual protium in the NMR solvent [$CHCl_3$, δ 7.26; $CHD_2OD$, δ 3.31; $(CHD_2)(CD_3)SO$, δ 2.49]. Data are represented as follows: chemical shift, multiplicity (s = singlet, d = doublet, t = triplet, q = quartet, quin = quintet, m = multiplet and/or multiple resonances, br = broad, app = apparent), integration, coupling constant in Hertz, and assignment. Proton-decoupled carbon nuclear magnetic resonance spectra ($^{13}C$ NMR) were recorded at 125 MHz at 24 °C, unless otherwise noted. Chemical shifts are expressed in parts per million (ppm, δ scale) downfield from tetramethylsilane and are referenced to the carbon resonances of the solvent ($CDCl_3$, δ 77.0; $CD_3OD$, δ 49.0; $(CD_3)_2SO$, δ 39.0). $^{13}C$ NMR and data are represented as follows: chemical shift, carbon type. Chemical shifts are expressed in parts per million (ppm, δ scale) downfield from tetramethylsilane. Infrared (IR) spectra were obtained using a Shimadzu 8400S FT-IR spectrometer referenced to a polystyrene standard. Data are represented as follows: frequency of absorption ($cm^{-1}$), intensity of absorption (s = strong, m = medium, w = weak, br = broad). High-resolution mass spectrometry (HRMS) measurements were obtained at the Chemistry and Chemical Biology Department, Harvard University Mass Spectrometry Facility using a Bruker microTOF-Q II hybrid quadrupole-time of flight, Agilent 1260 UPLC-MS. Low-resolution mass spectrometry (LRMS) measurements were obtained on Waters ACQUITY UPLC equipped with SQ Detector 2 mass spectrometer. Photo-irradiation was performed with a Dymax model 38100 UV curing light source flood lamp system with a ZIP shutter (Dymax, Torrington, CT). The absorbance was measured on a multi-mode microplate reader FilterMax F3 (Molecular Devices LLC, Sunnyvale, CA). Fluorescence and chemiluminescence signals were detected by scanning the gel on an Azure Imager C600 (Azure Biosystems, Inc., Dublin, CA).

**Experimental Procedures with Cell Lysates.**

**Preparation of homopropargylglycine labeled HEK 293T cell lysates.** HEK 293T cells were grown to confluency in a 15-cm plate in methionine-free media to which homopropargylglycine (1 mM) was added. The cells were grown for 24 h at 37 ˚C. The media was removed by aspiration and the cells were washed with PBS (2 × 10 mL) and harvested using trypsin. Cells were pelleted, the media was removed by aspiration, and the cell pellet was washed with PBS (2 × 10 mL). Cells were lysed with 1% RapiGest/PBS and EDTA-free protease inhibitors (1 mL) and sonicated on ice with a probe tip sonicator (12% power, 3 min). The cell lysate was cleared by centrifugation (21,130 × g) for 10 min at 4 ℃. The soluble protein concentration was determined using a BCA assay.

**CuAAC of homopropargylglycine-incorporated HEK 293T cell lysate with biotin-(PEG)$_3$-azide and the probes 5 and 6.** Based on BCA assay measurement, homopropargylglycine-incorporated HEK 293T cell lysate was adjusted to a protein concentration 1.3 mg/mL in 1% RapiGest/PBS. Cell lysate (100 µL) of was reacted with premixed click chemistry reagents (6.5 µL) at a final concentration of 100 µM cleavable biotin azide probe **5** or cleavable biotin picolyl azide probe **6**, 250 µM copper (II) sulfate, 250 µM THPTA, and 2.5 mM freshly prepared sodium ascorbate for 1.5 h at 24 ℃. The proteins were precipitated with methanol (400 µL) for 1 h at –80 ℃. The precipitated proteins were pelleted by centrifugation (21,130 g) for 10 min at 4 ℃. Methanol was discarded and the cell pellets were air dried for 15 min then resuspended in Laemmli sample buffer (15 µL). Results were analyzed by Western blot.

**Enrichment of homopropargylglycine-incorporated HEK 293T cell lysates for chemical proteomics.** HEK 293T cell lysates were adjusted to a protein concentration of 2.5 mg/mL in 1% RapiGest/PBS. For experiments in which CBPA optimized conditions were used for both probes **5** and **6**, cell lysates (500 µL) were reacted with pre-mixed click chemistry reagents (26 µL) at a final concentration of 100 µM of the cleavable biotin azide probe **5** or the cleavable biotin picolyl azide probe **6**, 250 µM copper(II)sulfate, 250 µM THPTA, and 2.5 mM freshly-prepared sodium ascorbate for 90 min at 24 ℃. For experiments in which individually optimized conditions for probe **5** were also used, cell lysates (500 µL) were reacted with pre-mixed click chemistry reagents (26 µL) at final concentrations of 200 µM of the cleavable biotin azide probe **5**, 300 µM copper(II)sulfate, 600 µM THPTA, and 2.5 mM freshly-prepared sodium ascorbate for 180 min at 24 ℃. The lysates were precipitated with methanol (1 mL) for 1 h at –80 ℃ then pelleted by centrifugation (21,130 × g) for 10 min at 4 ℃. Methanol was discarded and the cell pellets were air dried for 15 min. Biotinylated protein pellets were resuspended in 1% RapiGest/PBS (300 µL) and briefly sonicated with a probe tip sonicator. Streptavidin–agarose resin (200 µL 50% slurry, washed 3 × 1 mL PBS) was added to the resuspended protein solution and incubated for 12 h at 24 ℃ with rotation. The slurry was transferred to a filter column and the supernatant was removed by vacuum filtration. The beads were washed with 1% RapiGest/PBS (1 mL), urea (6 M, 2 × 1 mL), and PBS (2 × 1 mL). The washed beads were resuspended in PBS (200 µL), transferred to a 1.7 mL microcentrifuge tube, reduced with 5 mM dithiothreitol for 30 min at 24 ℃ with rotation, and alkylated with 10 mM iodoacetamide for 30 min at 24 ℃ in the dark with rotation. The beads were pelleted by centrifugation and the supernatant was removed. The beads were washed with PBS (1 mL) and resuspended in 0.5 M urea in PBS (200 µL). Trypsin (3 µL, 1.5 µg) was added and digestion was allowed to proceed for 12 h at 37 ℃ with rotation. The digest supernatant was collected and the beads were washed with PBS (200 µL) and water (2 × 200 µL). The washes were combined with the digest supernatant to afford the "trypsin fraction." The washed beads were treated with 2% formic acid/water (2 × 200 µL) for 30 min at 24 ℃. The supernatant was collected and the beads were washed with 1% formic acid/80% acetonitrile/water (2 × 200 µL). The washes were combined to afford the "cleavage fraction." Both the trypsin and cleavage fractions were concentrated to dryness using a SpeedVac concentrator and stored at –20 ℃ until analysis by LC-MS/MS. To desalt the samples, the samples were resuspended in 1% formic acid/water (50 µL) and loaded onto a C18 ZipTip. The loaded samples were washed with 1% formic acid/water (2 × 20 µL) and eluted with 1% formic acid/80%

acetonitrile/water (7 × 20 μL). The samples were concentrated to dryness using a SpeedVac concentrator and stored at –20 ℃ until analysis by LC-MS/MS. The desalted trypsin fractions were resuspended in TEAB buffer (50 mM, 20 μL). TMT reagent (36 mM, 2 μL) was added and allowed to incubate for 1 h at 24 ℃. 50% hydroxylamine/water (1 μL) was added to the TMT-labeled peptides and allowed to incubate for 15 min at 24 ℃ to quench excess TMT reagent. The TMT-labeled samples were combined and concentrated by SpeedVac.

**Experimental Procedures with Whole Cells.**

**Stimulation of A549 cells with phorbol 12-myristate 13-acetate (PMA).** A549 cells were grown to confluency in a 6-well plate and treated with DMEM containing 10 nM of PMA in DMSO. The cells were incubated for 6 h at 37 ℃. Media was removed by aspiration and the cells were washed with PBS (2 × 1 mL). The washed cells were then lysed directly in the plate with M-PER and EDTA-free protease inhibitor (200 µL). The lysate (180 µL) was collected and sonicated on ice with a probe tip sonicator (5 sec on, 2 sec off, 10% amplitude, 10 sec total). The sonicated cell lysate was cleared by centrifugation (21,130 × g) for 10 min at 4 ℃. The soluble protein concentration of each sample was determined using the BCA protein assay. Protein concentrations were adjusted to 2.03 mg/mL in M-PER. Protein expression was analyzed by Western blot.

**In-situ labeling of A549 cells with photo-celecoxib (2).** A549 cells were grown to confluency in a 15-cm diameter plate and then treated with FBS-free DMEM containing 10 µM of photo-celecoxib (**2**) or the minimalist tag **4** in DMSO. For competition experiments, 100 µM of celecoxib (**1**) in DMSO or 100 µM of licofelone (**3**) was added in addition to photo-celecoxib (**2**). The cells were incubated for 2 h at 37 ℃ then UV irradiated with a Dymax EC-5000 lamp for 2 min at 4 ℃. Irradiated cells were collected by scraping following trypsinization. Cells were pelleted and lysed with 1% RapiGest/PBS and EDTA-free protease inhibitor and sonicated on ice with a probe tip sonicator (2 sec on, 5 sec off, 10% amplitude, 10 sec total). Cell lysate was cleared by centrifugation (21,130 × g) for 10 min at 4 ℃ and the soluble protein concentration was determined by BCA assay. Protein concentrations were adjusted to 2.5 mg/mL in 1% RapiGest/PBS. Cell lysates were reacted with pre-mixed click chemistry reagents at final concentrations of 100 µM cleavable biotin picolyl azide probe **6**, 250 µM copper (II) sulfate, 250 µM THPTA, and 2.5 mM freshly prepared sodium ascorbate for 90 min at 24 ℃. The proteins were precipitated with methanol for 1 h at –80 ℃ then pelleted by centrifugation (21,130 × g) for 10 min at 4 ℃. Methanol was discarded and protein pellets were air dried for 15 min prior to analysis by Western blot. Proteins were quantified using ImageStudioLite v5.2.5.

**Enrichment of photo-celecoxib-conjugated proteins for LC-MS/MS.** Biotinylated protein pellets were resuspended in 1% RapiGest/PBS (300 µL) and briefly sonicated. Streptavidin–agarose resin (200 µL of 50% slurry, washed 3 × 1 mL PBS) was added to the resuspended protein solution and incubated for 12 h at 24 ℃ with rotation. The slurry was transferred to a filter column and the supernatant was removed by vacuum filtration. The beads were washed with 1% RapiGest/PBS (1 mL), urea (6M, 2 × 1 mL), and PBS (2 × 1 mL). The washed beads were resuspended in PBS (200 µL) then reduced with 5 mM dithiothreitol (DTT) for 30 min at 24 ℃ and alkylated with 10 mM iodoacetamide for 30 min at 24 ℃ in the dark with rotation. The beads were pelleted by centrifugation and the supernatant was removed. The beads were washed with PBS (1 mL) and resuspended in 0.5 M urea in PBS (200 µL). Trypsin (1.5 µg) was added to the samples and digestion was allowed to proceed for 12 h at 37 ℃ with rotation. The digest supernatant was collected and the beads were washed with PBS (200 µL) and water (2 × 200 µL). The washes were combined to afford the "trypsin fraction." The probe **6** was cleaved to recover the conjugated peptide in 2% formic acid/water (2 × 200 µL) for 30 minutes at 24 ℃. The supernatant was collected and the beads were washed with 80% acetonitrile and 1% formic acid/water (2 × 200 µL). The washes were combined to afford the "cleavage fraction." Both the trypsin and cleavage fractions were concentrated to dryness using a SpeedVac concentrator. To desalt the samples, the samples were resuspended in 1% formic acid/water (50 µL) and loaded onto a C18 ZipTip. The loaded samples were washed with 1% formic acid/water (2 × 20 µL) and eluted with 1% formic acid/80% acetonitrile/water (7 × 20 µL). The samples were concentrated to dryness using a SpeedVac concentrator and stored at –20 ℃ until analysis by LC-MS/MS. The desalted trypsin fractions were resuspended in TEAB buffer (50 mM, 20 µL). TMT reagent (36 mM, 2.0 µL) was added and incubated for 1 h at 24 ℃. 50% hydroxylamine/water (1 µL) was added and allowed to incubate for 15 min at 24 ℃ to quench excess TMT reagent. The TMT-labeled samples were combined and concentrated by SpeedVac.

**Mass Spectrometry Procedures.**

The desalted samples were resuspended in 0.1% formic acid/water (20 µL). The sample (10.0 µL) was loaded onto a C18 trap column (3 cm, 3 µm particle size C10 Dr. Maisch 150 µm I.D) and then separated on an analytical column (Thermo Scientific Acclaim PepMap 100, 2 µm particle size, 250 mm length, 75 µm internal diameter) at 150 nL/min with a Thermo Scientific Ultimate 3000 system connected in line to a Thermo Scientific Orbitrap Fusion Tribrid. The column temperature was maintained at 50 ˚C. The tryptic peptides were separated via a step-wise gradient from 5% to 98% of 0.1% formic acid/acetonitrile over 120 min (0–1 min, 0–5%; 1–91 min, 5–27%; 91–115 min, 27–98%; 115–120 min, 98%–0%). The cleavage peptides were separated via the same gradient described above. Survey scans of peptide precursors were performed at 120K FWHM resolution ($m/z$ = 200). Tandem MS was performed on the most abundant precursors exhibiting a charge state from 2 to 6 at a resolving power settings of 50K. HCD fragmentation was applied with 35% collision energy and resulting fragments accumulated for up to 100 ms.

**Data Analysis Procedures.**

**Quantitative interactome analysis.** Analysis was performed in Thermo Scientific Proteome Discoverer version 2.3. HCD spectra with a signal-to-noise ratio greater than 1.5 were searched against a database containing the Uniprot 2016 annotated human proteome and contaminant proteins using Sequest HT with a mass tolerance of 10 ppm for the precursor and 0.02 Da for fragment ions with specific trypsin digestion, 2 missed cleavages, variable oxidation on methionine residues (+15.995 Da), static carboxyamidomethylation of cysteine residues (+57.021 Da), and static TMT labeling at lysine residues and N-termini. Assignments were validated using Percolator. The resulting assignments were filtered to only include high-confidence matches, and TMT reporter ions were quantified using the Reporter Ions Quantifier. Only non-contaminant proteins with at least 3 unique peptides and high confidence FDR were considered for interactome analysis.

**Quantitative analysis of the cleavage fraction of HPG-labeled lysates.** For HPG-labeled samples, SEQUEST HT results from CBA and CBPA samples were aligned with differences in retention times approximated by a linear model. The alignment was used to predict the retention time of peptides unassigned in either of the two samples. PSMs were then quantified by peak intensities. The code used for this analysis is available on github: https://github.com/christinawoo/CBPA_CPA The intensities of peptides detected with both handles were used to evaluate the improvement in enrichment.

**Analysis of the cleavage fraction following photo-celecoxib enrichment.** Data analysis was performed with Proteome Discoverer version 2.3 using SEQUEST HT, allowing for variable modifications (methionine oxidation: +15.995 Da; cysteine carbamidomethylation: +57.021 Da; photo-celecoxib with CBPA: +620.177 Da, +622.184 Da), up to two missed cleavages and a mass tolerance of 10 ppm for the precursor ion and 0.02 Da for fragment ions from HCD. Searching was performed against the Swiss-Prot human database and a contaminant protein database. For binding sites of photo-celecoxib, MS/MS data from the cleavage fraction were searched by SEQUEST HT against the focused protein list observed from the trypsin fraction. The high confidence peptide assignments (false discovery rate < 1%) were analyzed using IsoStamp[8] for the precursor isotope pattern and filtered based on manual validation.

**Structural Modeling Procedures.**

Photo-celecoxib (**2**) was minimized using Gaussian 16 with the HF/6-31g(d) basis set. The minimized photo-celecoxib (**2**) was added to the structure of PTGES using PDB structure 4YK5 as a model. Simulated annealing was then performed using Amber 18. The ff14SB forcefield was used for the protein, and the GAFF-derived forcefield was used for photo-celecoxib (**2**). The system was solvated with an 8 Å box of TIP3P water atoms. The backbone of the protein was restrained to the starting position with force constant 5. Restraints were placed on the distance between the diazirine carbon and nitrogen atoms and the protonated nitrogen atom of the side chain of the labeled histidine residue. The restraint between the diazirine carbon atom and the histidine nitrogen atom had a minimum distance of 5 Å and a force constant of 5. The restraints between the diazirine nitrogen atoms and the histidine nitrogen atom had a minimum distance of 3 Å and a force constant of 5. The system was heated from 0 ºK to 800 ºK over 1,000 steps, held at 800 ºK for 2,000 steps, and cooled to 0 ºK over 12,000 steps, with a step time of 0.002 ps. This process was repeated ten times from the same starting structure. Of the ten structures, two showed geometries that would allow for the observed labeling. The lowest energy of these two structures was then further minimized by heating to 300 ºK over 1,000 steps, then holding at 300 ºK for 4,000 steps, with a step time of 0.002 ps. During this optimization, the backbone of the protein was restrained to its starting position with a force constant of 5, and the force constants of the restraints on the diazirine carbon and nitrogen atoms and the protonated nitrogen atom of the side chain of the labeled histidine residue were increased to 25.

**Synthetic Procedures.**

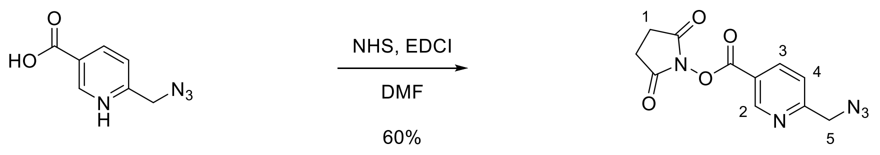


**S1** **S2**

***Synthesis of 2,5-dioxopyrrolidin-1-yl 6-(azidomethyl)nicotinate (S2):***

*N*-(3-Dimethylaminopropyl)-*N'*-ethylcarbodiimide hydrochloride (1.32 g, 6.89 mmol, 1.50 equiv) was added to a solution of the pyridine **S1** (819 mg, 4.60 mmol, 1 equiv) in dimethylformamide (23 mL) and the resulting solution was stirred for 5 min at 24 ˚C. *N*-Hydroxysuccinimide was added and the resulting solution was stirred for 18 h at 24 ˚C. The product mixture was concentrated by rotary evaporation and the residue obtained was purified by flash column chromatography ($SiO_2$, 0–5% methanol–dichloromethane, 3 steps) to afford the NHS ester **S2** as a light orange solid (754 mg, 60%).

$R_f$ = 0.52 (5% methanol/DCM, UV). $^1$H NMR (500 MHz, $CDCl_3$): δ 9.25 (d, 1H, $J$ = 2.1 Hz, $H_2$), 8.39 (dd, 1H, $J$ = 8.2, 2.2 Hz, $H_3$), 7.52 (d, 1H, $J$ = 8.2 Hz, $H_4$), 4.60 (s, 2H, $H_5$), 2.90 (s, 4H, $H_1$). $^{13}$C NMR (125 MHz, $CDCl_3$): δ 169.0 (C), 162.3 (C), 160.6 (2 × C), 151.3 (CH), 139.0 (CH), 121.6 (CH), 120.8 (C), 55.3 ($CH_2$), 25.7 (2 × $CH_2$). IR (ATR-FTIR), $cm^{-1}$: 2103 (s), 1734 (s), 1597 (m), 1198 (m), 1067 (m), 724 (s). HRMS-ESI (*m/z*): $[M+H]^+$ calculated for $C_{11}H_{10}N_5O_4$, 276.0727; found, 276.0732.

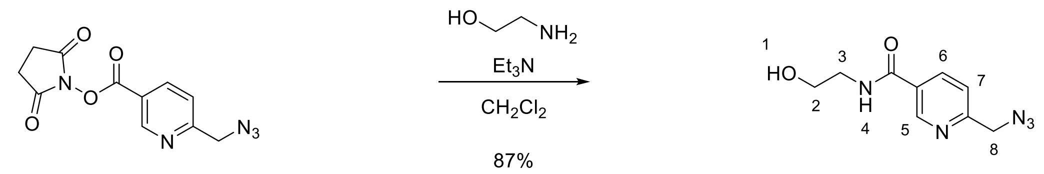


***Synthesis of 6-(azidomethyl)-N-(2-hydroxyethyl)nicotinamide (S3):***

Triethylamine (350 µL, 1.33 mmol, 1.50 equiv) and ethanolamine (80.0 µL, 1.33 mmol, 1.50 equiv) were added sequentially to a solution of the NHS ester **S2** (244 mg, 0.89 mmol, 1 equiv) in dichloromethane (7.0 mL) and the resulting solution was stirred for 18 h at 24 °C. The product mixture was diluted with brine (15 mL), and the biphasic mixture was transferred to a separatory funnel. The layers that formed were separated. The aqueous layer was extracted with 10% isopropanol–ethyl acetate (3 × 15 mL), and the organic layers were combined. The combined organic layers were dried over sodium sulfate. The dried solution was filtered and the filtrate was concentrated by rotary evaporation. The residue obtained was purified by flash column chromatography ($SiO_2$, 2–5% methanol–dichloromethane, 2 steps) to afford the amide **S3** as an off-white solid (171 mg, 87%).

$R_f$ = 0.32 (5% methanol–dichloromethane, UV). $^1$H NMR (500 MHz, $CDCl_3$): δ 8.91 (s, 1H, $H_5$), 8.09 (d, 1H, *J* = 8.1 Hz, $H_6$), 7.66 (t, 1H, *J* = 5.1 Hz, $H_4$), 7.32 (d, 1H, *J* = 8.1 Hz, $H_7$), 4.45 (s, 2H, $H_8$), 4.38 (bs, 1H, $H_1$), 3.77–3.75 (m, 2H, $H_2$), 3.56–3.53 (m, 2H, $H_3$). $^{13}$C NMR (125 MHz, $CDCl_3$): δ 166.2 (C), 158.6 (C), 148.0 (CH), 136.5 (CH), 129.3 (C), 121.8 (CH), 61.3 ($CH_2$), 55.1 ($CH_2$), 42.9 ($CH_2$). IR (ATR-FTIR), $cm^{-1}$: 3293 (br), 2098 (s), 1637(s), 1295 (m), 1063 (m). HRMS-ESI (*m/z*): $[M+H]^+$ calculated for $C_9H_{12}N_5O_2$, 222.0986; found, 222.1006.

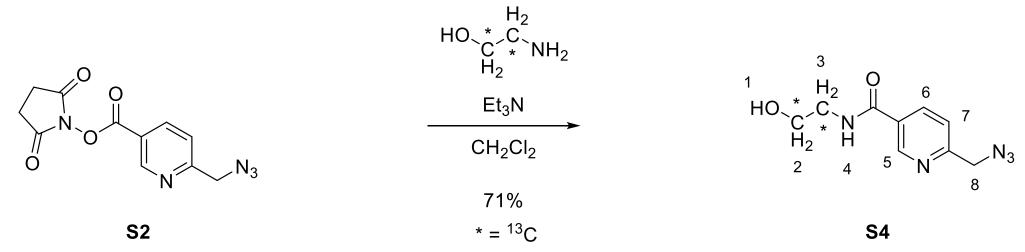

*Synthesis of 6-(azidomethyl)-N-(2-hydroxyethyl-1,2-13C2)nicotinamide (S4):*

Ethanolamine-$^{13}C_2$ hydrochloride (99.0 mg, 0.99 mmol, 1.30 equiv) and triethylamine (401 µL, 1.50 mmol, 2.00 equiv) were added sequentially to a solution of the NHS ester **S2** (210 mg, 800 µmol, 1 equiv) in dichloromethane (7.6 mL) and the resulting solution was stirred for 18 h at 24 °C. The product mixture was diluted with brine (15 mL), and the biphasic mixture was transferred to a separatory funnel. The layers that formed were separated. The aqueous layer was extracted with 10% isopropanol–ethyl acetate (3 × 15 mL), and the organic layers were combined. The combined organic layers were dried over sodium sulfate. The dried solution was filtered and the filtrate was concentrated by rotary evaporation. The residue obtained was purified by flash column chromatography ($SiO_2$, 2–5% methanol–dichloromethane, 2 steps) to afford the amide-$^{13}C_2$ **S4** as an off-white solid (122 mg, 71%).

$R_f$ = 0.40 (5% methanol–dichloromethane, UV). $^1$H NMR (500 MHz, $CDCl_3$): δ 8.92 (s, 1H, $H_5$), 8.09 (d, 1H, $J$ = 8.1 Hz, $H_6$), 7.68 (bs, 1H, $H_4$), 7.32 (d, 1H, $J$ = 8.1 Hz, $H_7$), 4.45 (s, 2H, $H_8$), 4.41 (bs, 1H, $H_1$), 3.91–3.88 (m, 1H, $H_2$), 3.69–3.65 (m, 1H, $H_2$), 3.61 (bs, 1H, $H_3$), 3.42–3.38 (m, 1H, $H_3$). $^{13}$C NMR (150 MHz, $CDCl_3$): δ 166.2 (C), 158.6 (C), 148.0 (CH), 136.5 (CH), 129.3 (C), 121.8 (CH), 61.3 (d, $J_{cc}$ = 38.2 Hz, $^{13}CH_2$), 55.1 ($CH_2$), 42.9 (d, $J_{cc}$ = 38.5 Hz, $^{13}CH_2$). IR (ATR-FTIR), cm$^{-1}$: 3394 (br), 2930 (m), 1698 (m), 1253 (m), 828 (s). HRMS-ESI (*m/z*): $[M+H]^+$ calculated for $C_7{}^{13}C_2H_{12}N_5O_2$, 224.1058; found, 224.1080.

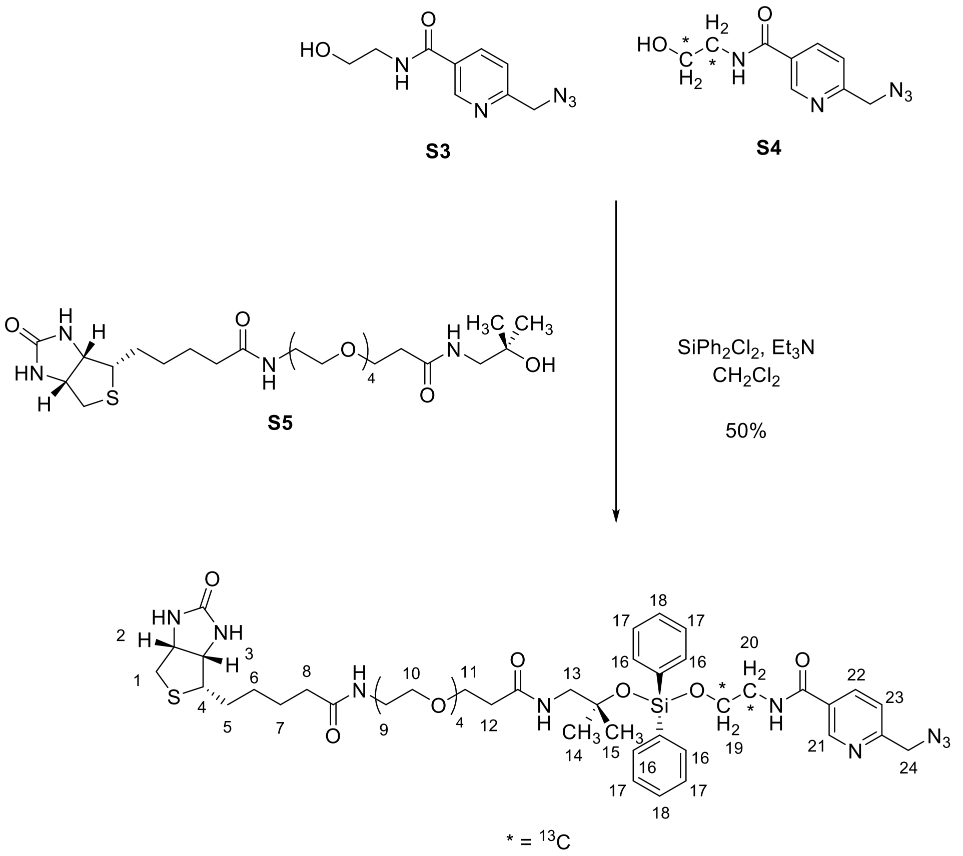

cleavable biotin picolyl azide probe **6** ($^{13}$C, 75%)

***Synthesis of the cleavable biotin picolyl azide probe 6:***

Triethylamine (55.0 µL, 400 µmol, 20.0 equiv) and dichlorodiphenylsilane (10.7 µL, 51.0 µmol, 3.00 equiv) were added in sequence to a solution of biotin-CA(PEG)$_4$-alcohol **S5** (11.1 mg, 19.1 µmol, 1 equiv) in dichloromethane (99 µL) and the resulting solution was stirred for 2 h at 24 ˚C. A 1:3 mixture of the amide **S3** and the amide-$^{13}C_2$ **S4** (26.0 mg, 118 µmol, 6.00 equiv) in dichloromethane (400 µL) was added and the solution was stirred for 18 h at 24 ˚C. The reaction mixture was diluted with aqueous sodium bicarbonate (1 mL) and the layers that formed were separated. The aqueous layer was extracted with dichloromethane (3 × 1 mL), and the organic layers were combined. The combined organic layers were dried over sodium sulfate. The dried solution was filtered and the filtrate was concentrated by rotary evaporation. The residue obtained was purified by flash column chromatography ($SiO_2$, 1–11% methanol-dichloromethane, 10 steps) to afford the cleavable biotin picolyl azide probe **6** as a yellow oil (9.61 mg, 50%).

$R_f$ = 0.30 (10% methanol–dichloromethane, $I_2$). $^1$H NMR (600 MHz, $CD_3OD$): δ 8.89 (d, 1H, $J$ = 2.0 Hz, $H_{21}$), 8.16 (dd, 1H, $J$ = 8.1, 2.3 Hz, $H_{22}$), 7.61 (d, 4H, $J$ = 6.8 Hz, $H_{16}$), 7.53 (d, 1H, $J$ = 8.2 Hz, $H_{23}$), 7.37 (t, 2H, $J$ = 7.4 Hz, $H_{18}$), 7.30 (t, 4H, $J$ = 7.4 Hz, $H_{17}$), 4.55 (s, 2H, $H_{24}$), 4.45 (dd, 1H, $J$ = 7.9, 4.9 Hz, $H_2$), 4.27 (dd, 1H, $J$ = 4.5, 7.8 Hz, $H_3$), 4.08–4.05 (m, 1H, $H_{19}$), 3.95 (t, 2H, $J$ = 5.7 Hz, $H_{19}$), 3.84–3.81 (m, 1H, $H_{19}$), 3.71–3.69 (m, 1H, $H_{20}$), 3.68 (t, 2H, $J$ = 6.1 Hz, $H_{11}$), 3.60–3.52 (m, 15H, $H_9$/$H_{10}$/$H_{20}$), 3.50 (t, 2H, $J$ = 5.5 Hz, $H_{13}$), 3.48–3.46 (m, 1H, $H_{20}$), 3.32 (t, 2H, $J$ = 5.4 Hz, $H_9$), 3.17 (dt, 1H, $J$ = 8.5, 5.4 Hz, $H_4$), 2.89 (dd, 1H, $J$ = 12.7, 5.0 Hz, $H_1$), 2.67 (d, 1H, $J$ = 12.7 Hz, $H_1$), 2.44 (t, 2H, $J$ = 6.1 Hz, $H_{12}$), 2.19 (t, 2H, $J$ = 7.4 Hz, $H_8$), 1.74–1.52 (m, 4H, $H_5$/$H_7$), 1.41 (quin, 2H, $J$ = 7.6 Hz, $H_6$), 1.22 (s, 6H, $H_{14}$/$H_{15}$). $^{13}$C NMR

(150 MHz, $CD_3OD$): δ 176.1 (C), 174.1 (C), 167.7 (C), 166.1 (C), 160.0 (C), 149.4 (CH), 137.8 (CH), 136.1 (4 × CH), 135.5 (2 × C), 131.3 (2 × CH), 130.9 (C), 128.9 (4 × CH), 123.3 (CH), 77.0 ($CH_2$), 71.6 ($CH_2$), 71.5 ($CH_2$), 71.5 ($CH_2$), 71.5 ($CH_2$), 71.3 ($CH_2$), 71.3 ($CH_2$), 70.6 ($CH_2$), 68.4 ($CH_2$), 63.4 (CH), 62.7 (d, $J$ = 48.3 Hz, $^{13}CH_2$), 61.6 (CH), 61.3 (CH), 57.0 (CH), 55.9 ($CH_2$), 51.6 (C), 43.2 (d, $J$ = 48.2 Hz, $^{13}CH_2$), 41.1 ($CH_2$), 40.4 ($CH_2$), 37.8 ($CH_2$), 36.7 ($CH_2$), 29.8 ($CH_2$), 29.5 ($CH_2$), 28.2 (2 × $CH_3$), 26.8 ($CH_2$). IR (ATR-FTIR), $cm^{-1}$: 3306 (br), 2926 (m), 2101 (s), 1643 (s), 1116 (s). HRMS-ESI (*m/z*): $[M+Na]^+$ calculated for $C_{46}H_{65}N_9O_{10}SSiNa$/$C_{44}{}^{13}C_2H_{65}N_9O_{10}SSiNa$, 986.4237/988.4303; found, 986.4185/988.4287.

**Catalog of Nuclear Magnetic Resonance and Infrared Spectra.**

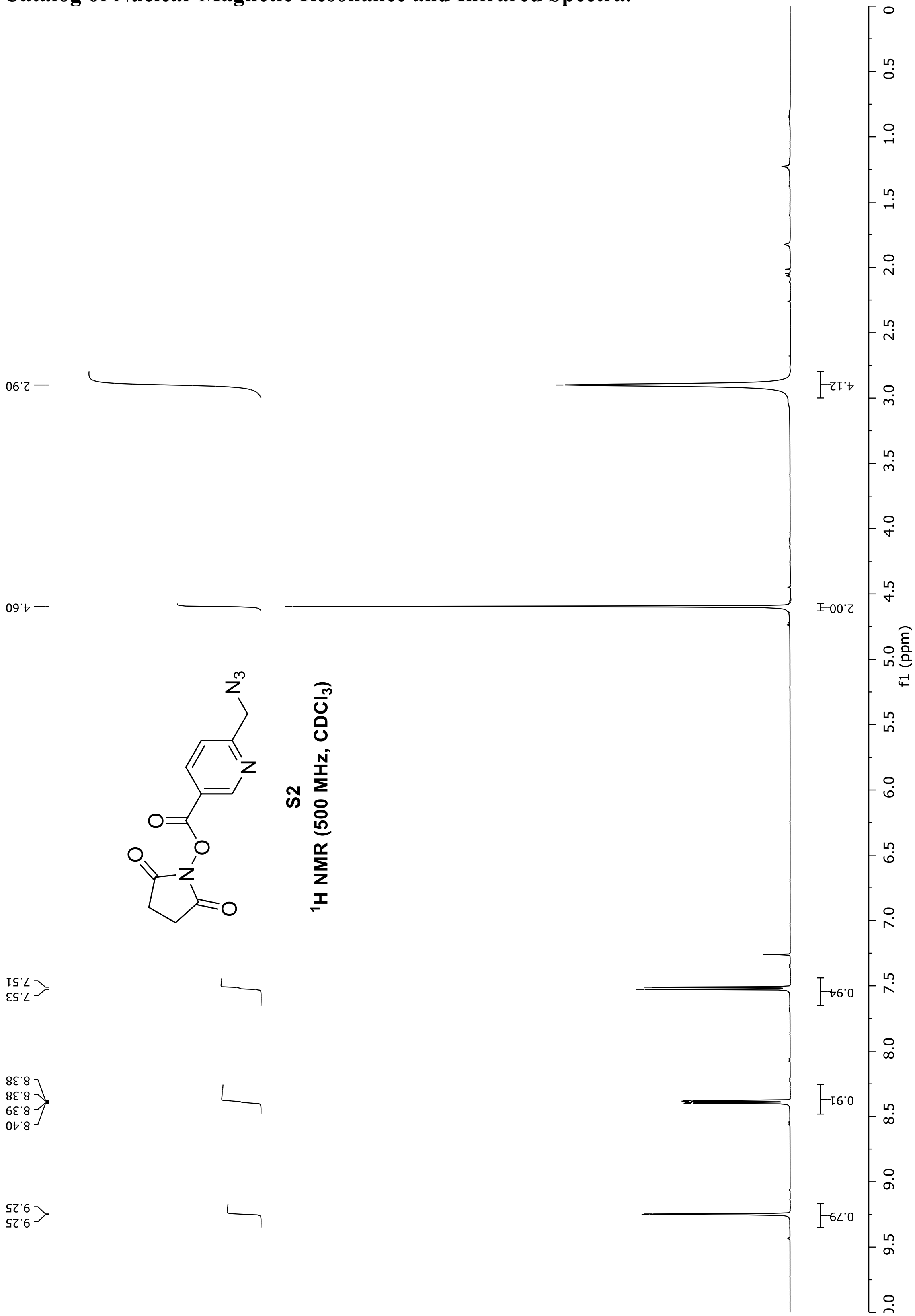

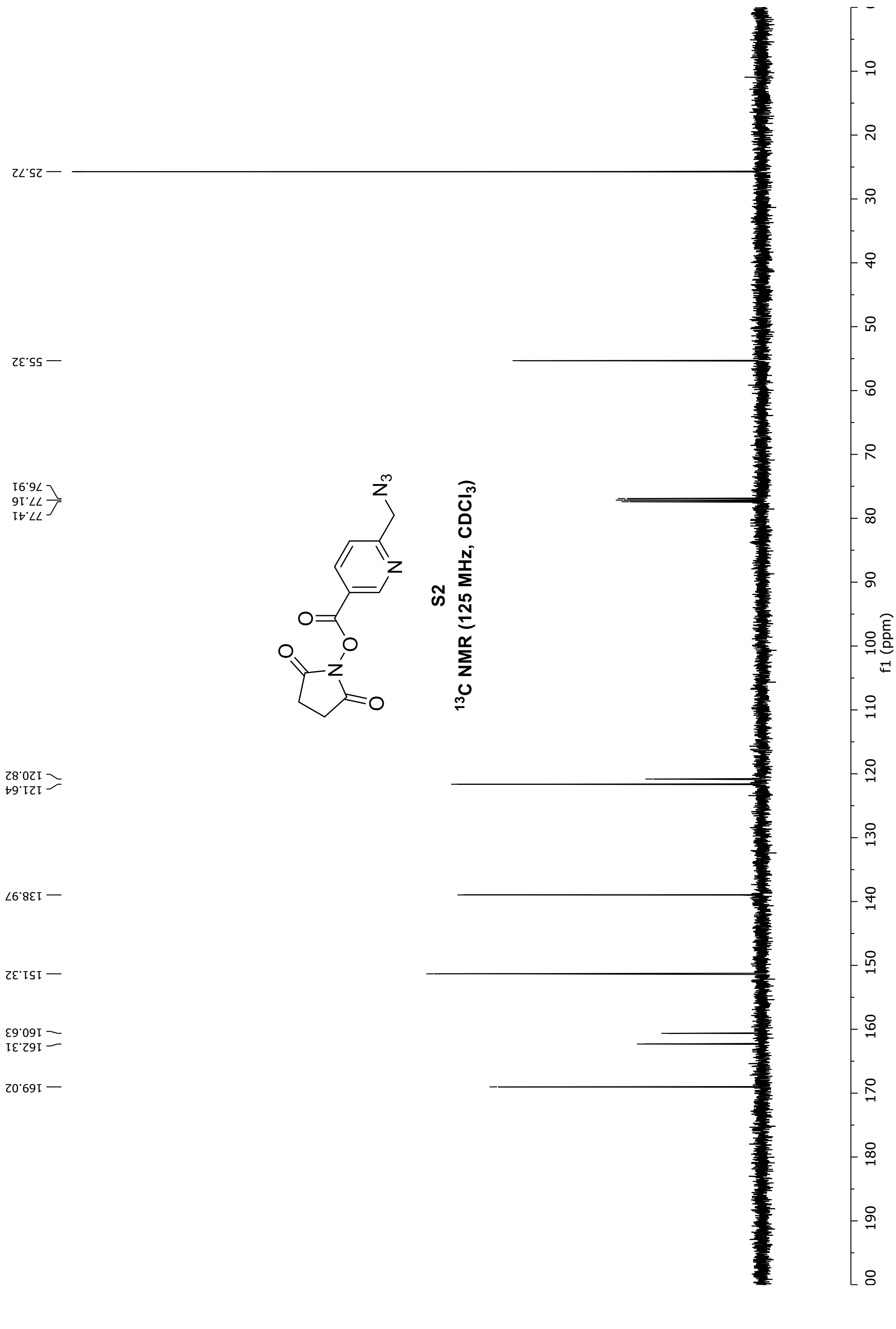


Miyamoto, D.; et al. “Discovery of a celecoxib binding site on PTGES with a cleavable chelation-assisted biotin probe” *ACS Chem. Biol.* **2019**.

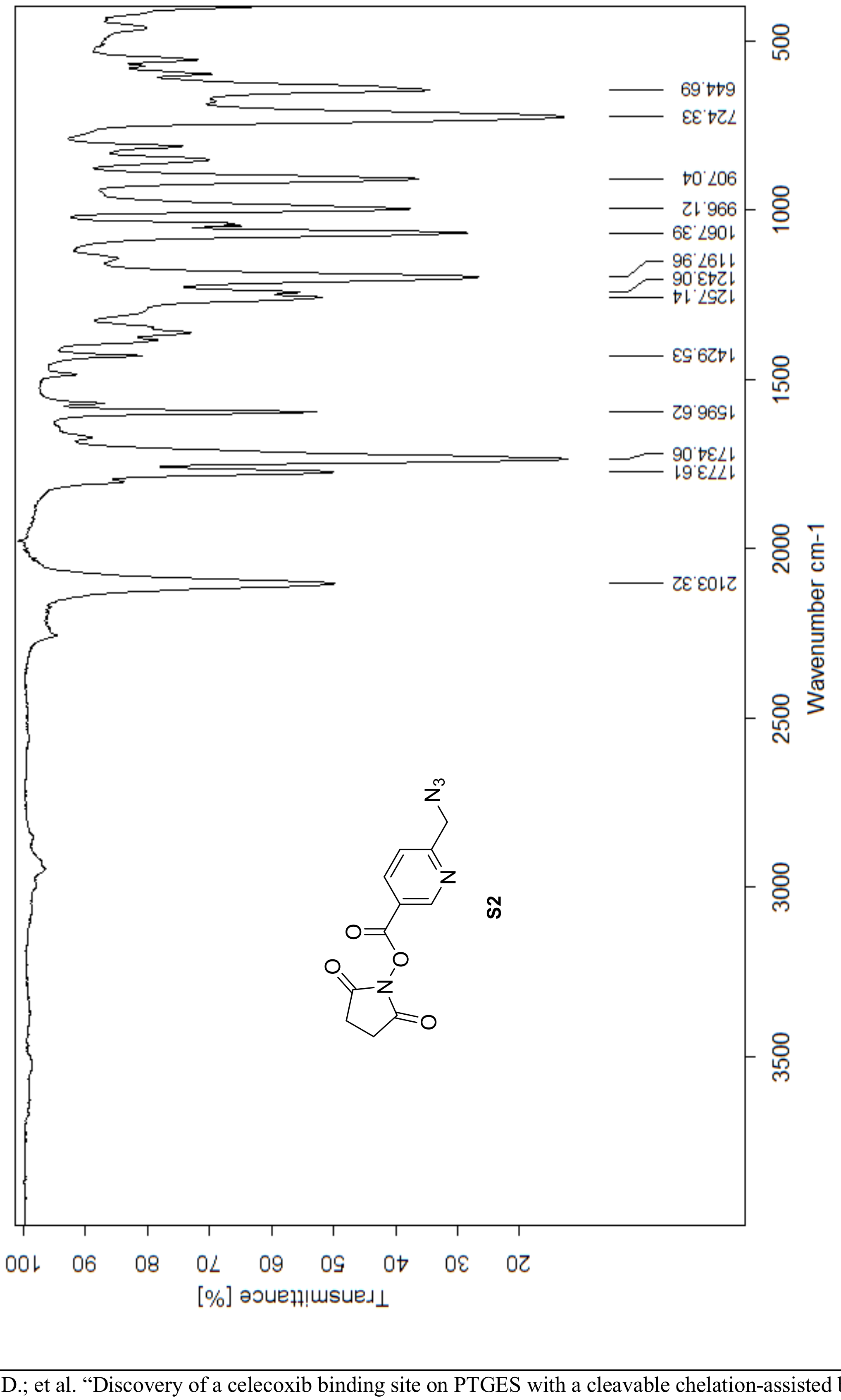

Transmittance [%]
Wavenumber cm-1
644.69
724.33
907.04
996.12
1067.39
1197.96
1243.06
1257.14
1429.53
1596.62
1734.06
1773.61
2103.32
S2

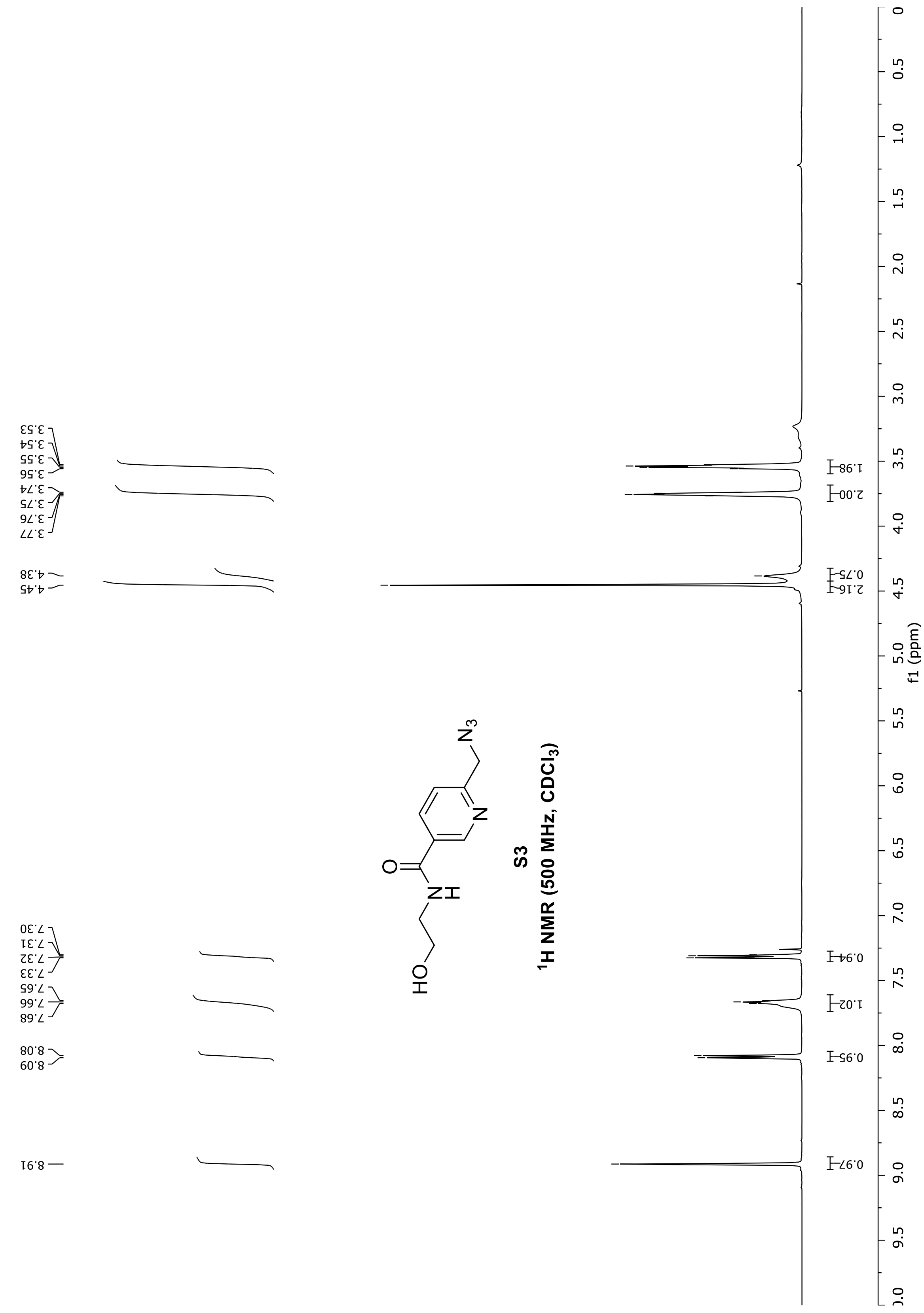
S3
1H NMR (500 MHz, CDCl3)
f1 (ppm)

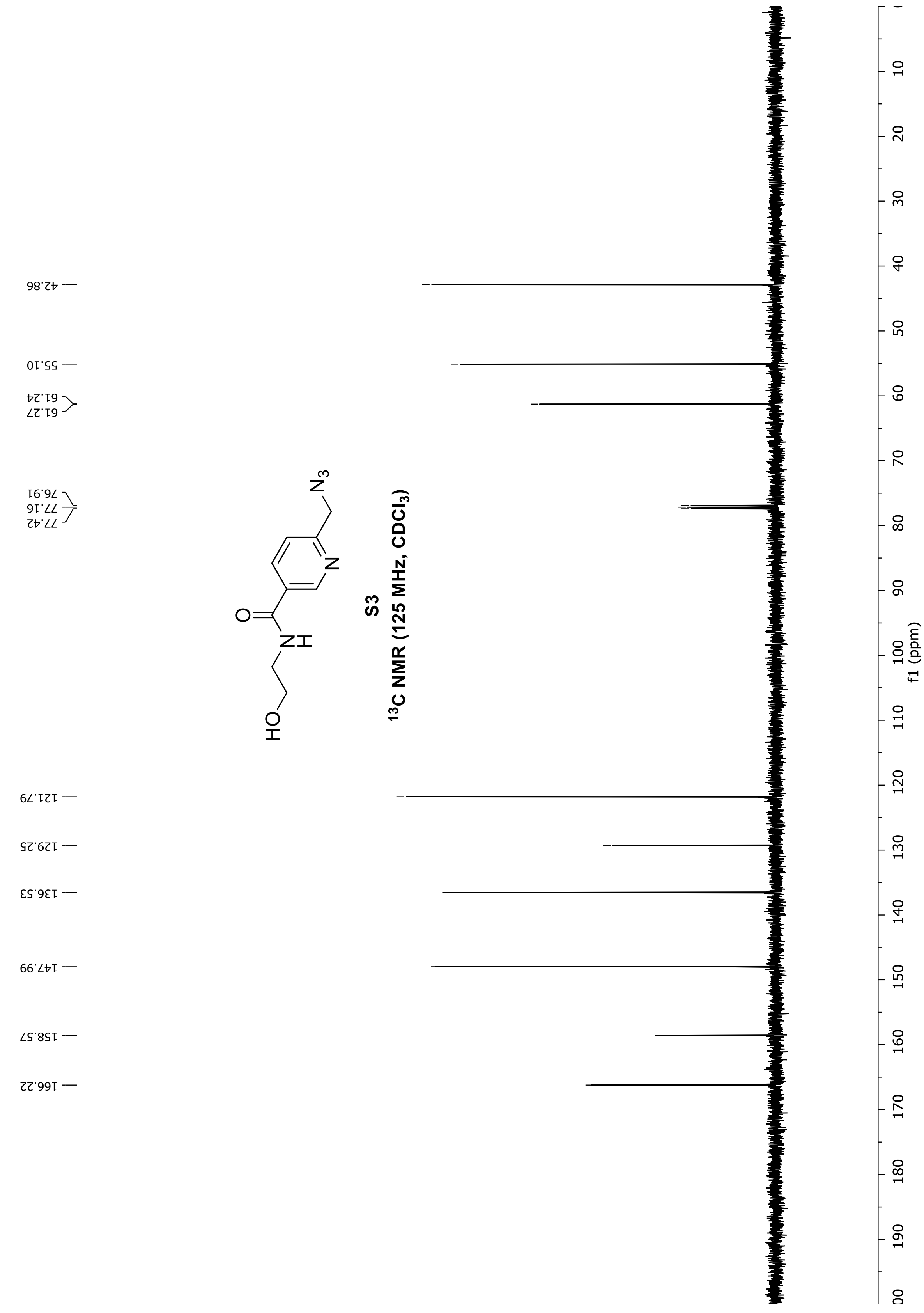
42.86
55.10
61.24
61.27
76.91
77.16
77.42
121.79
129.25
136.53
147.99
158.57
166.22
S3
13C NMR (125 MHz, CDCl3)
f1 (ppm)

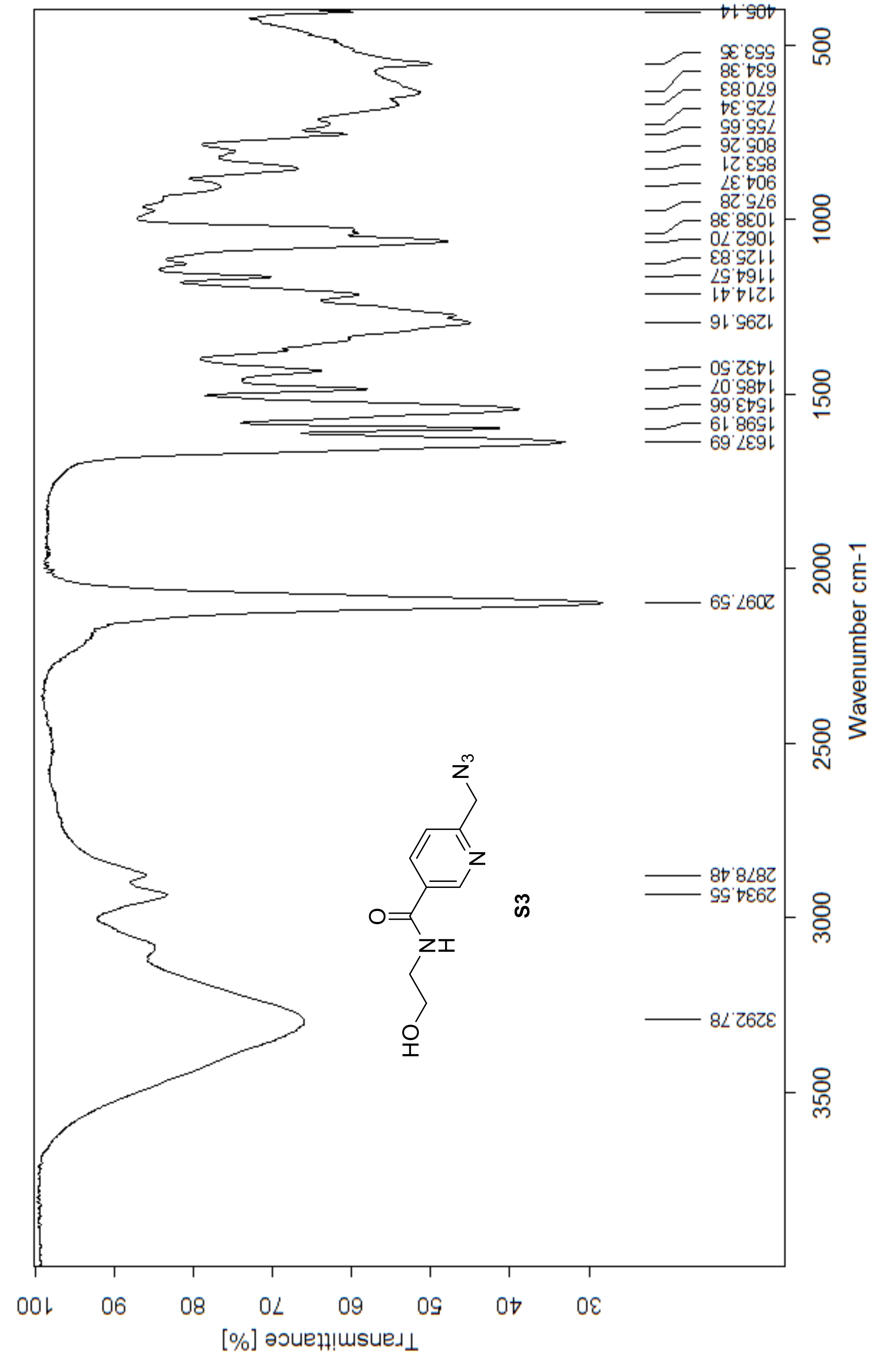


Miyamoto, D.; et al. "Discovery of a celecoxib binding site on PTGES with a cleavable chelation-assisted biotin probe" *ACS Chem. Biol.* **2019**.

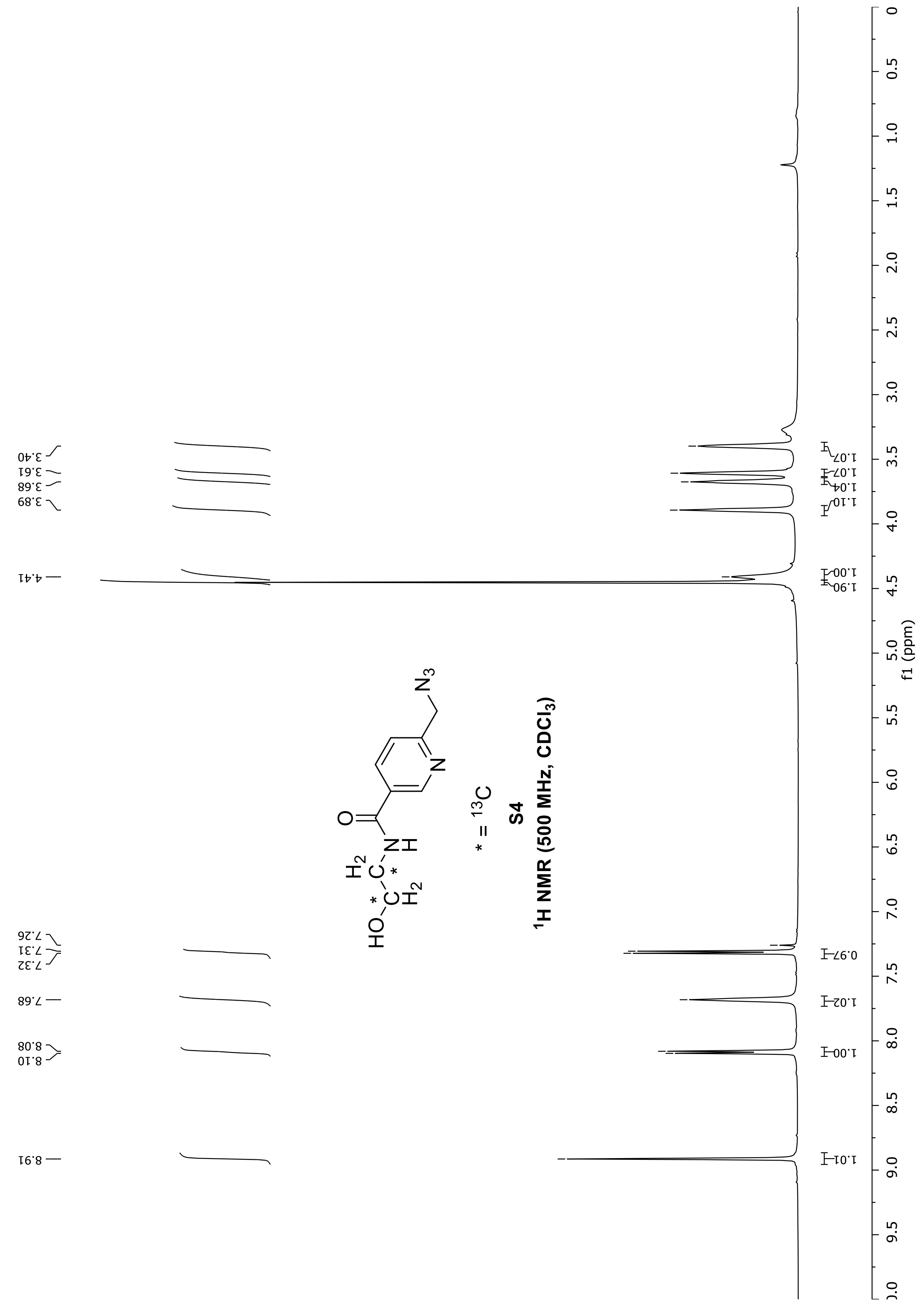

* = $^{13}C$
S4
$^1H$ NMR (500 MHz, $CDCl_3$)
f1 (ppm)

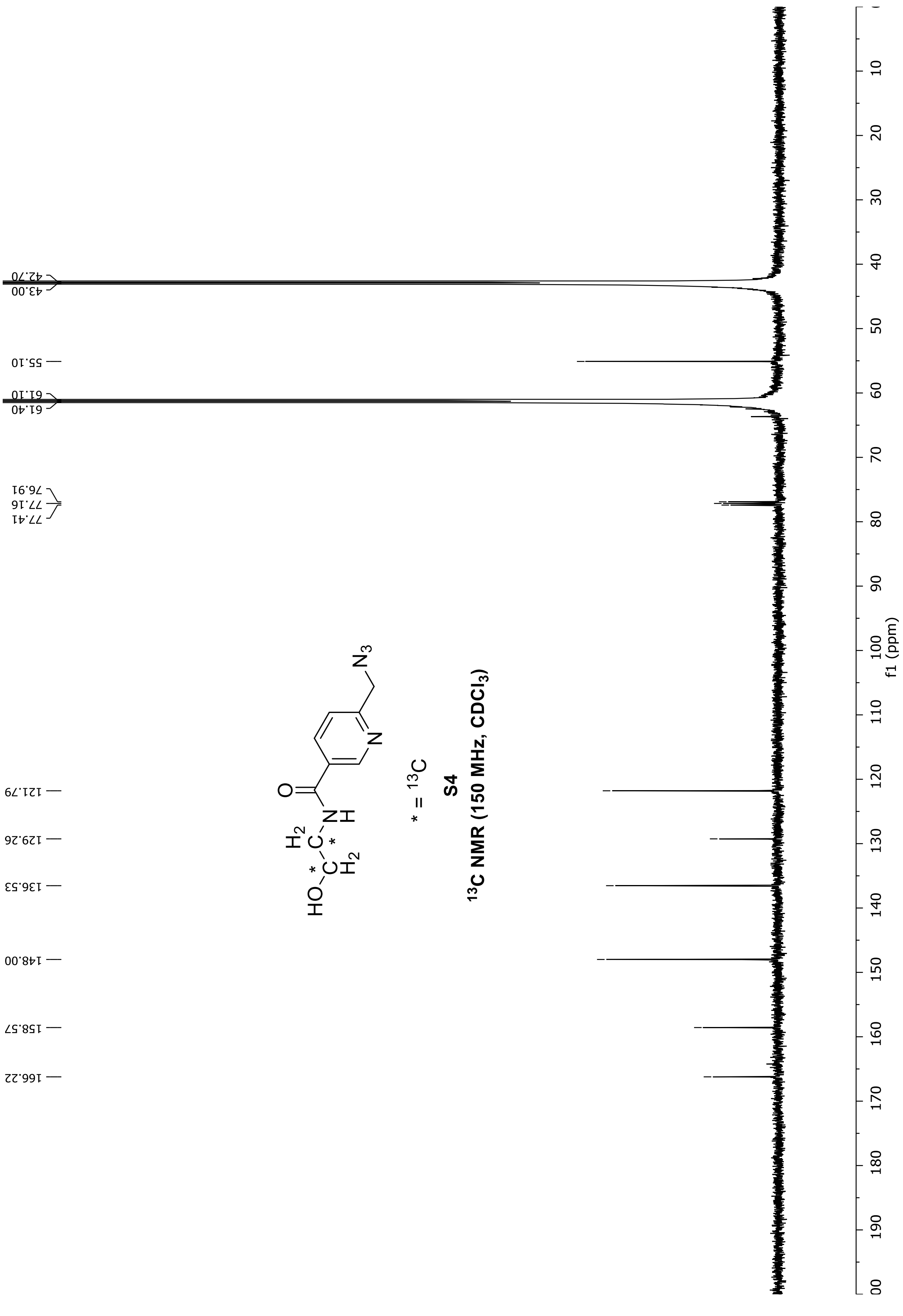

42.70
43.00
55.10
61.10
61.40
76.91
77.16
77.41
121.79
129.26
136.53
148.00
158.57
166.22
N3
N
O
H
N
H2
C
C
H2
HO
OH
* = 13C
S4
13C NMR (150 MHz, CDCl3)
f1 (ppm)
0
10
20
30
40
50
60
70
80
90
100
110
120
130
140
150
160
170
180
190
200

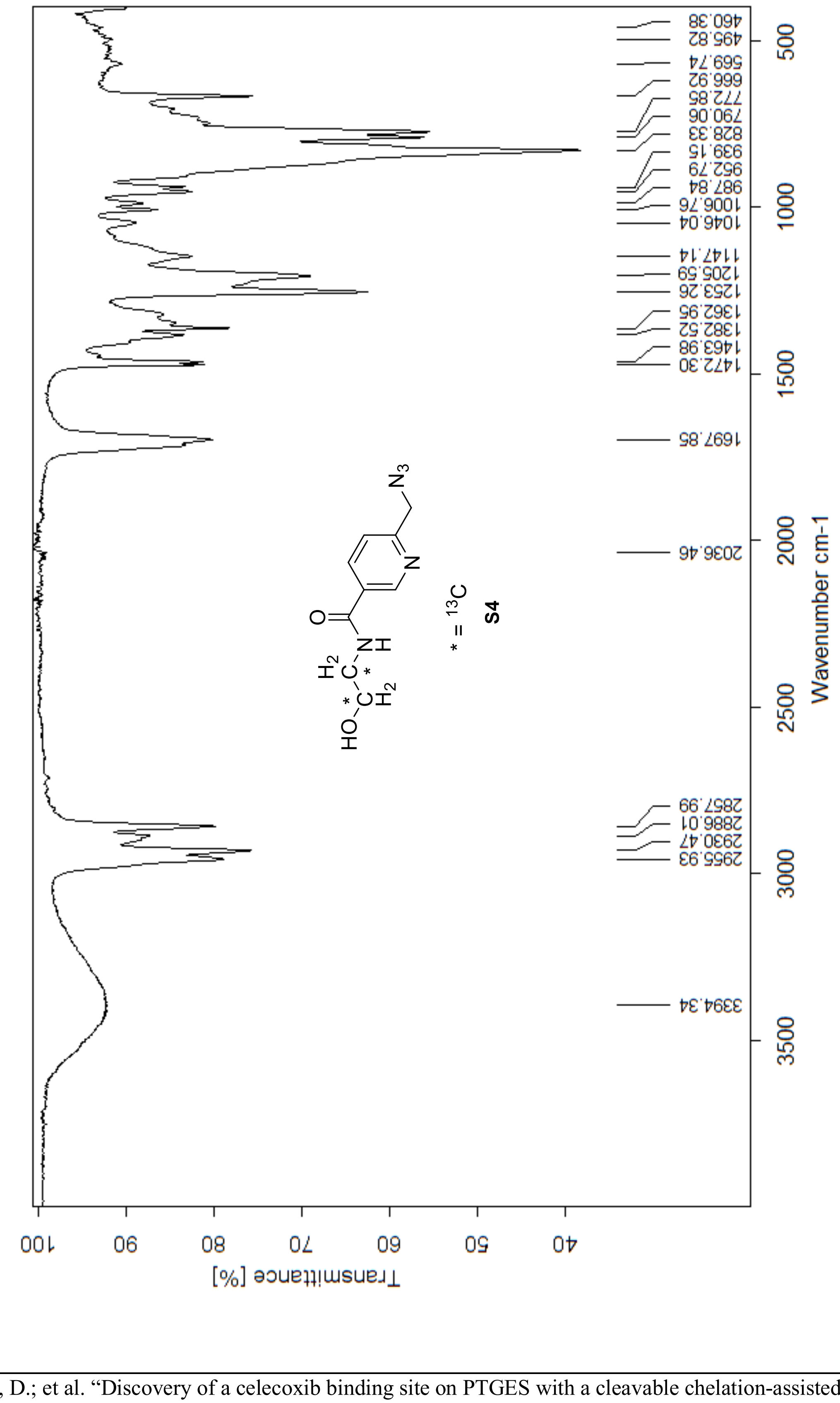
Transmittance [%]
Wavenumber cm-1
460.38
495.82
569.74
666.92
772.85
790.06
828.33
939.15
952.79
987.84
1006.76
1046.04
1147.14
1205.59
1253.26
1362.95
1382.52
1463.98
1472.30
1697.85
2036.46
2857.99
2886.01
2930.47
2955.93
3394.34
* = 13C
S4

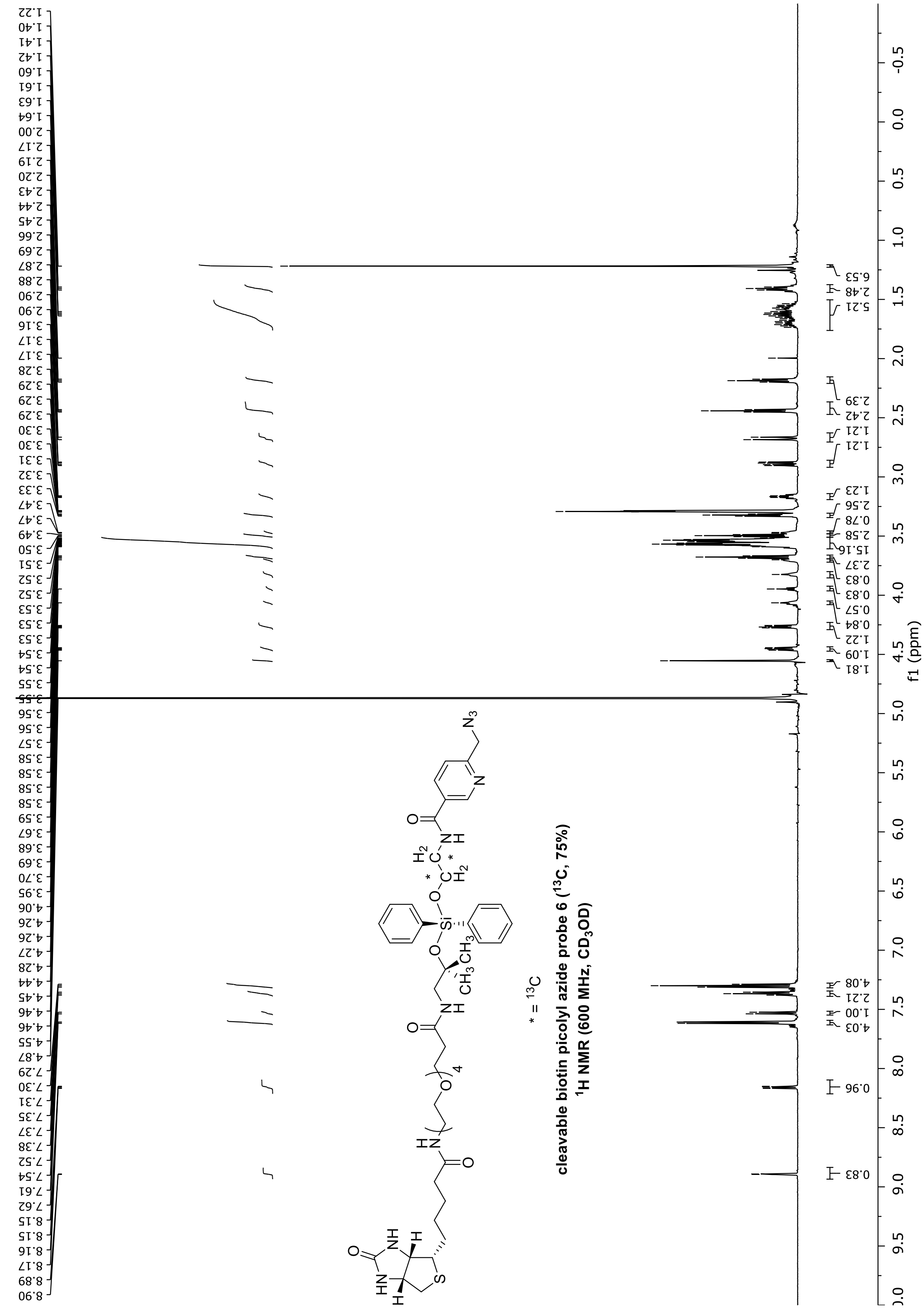
cleavable biotin picolyl azide probe 6 ($^{13}C$, 75%)
$^1H$ NMR (600 MHz, $CD_3OD$)
* = $^{13}C$
f1 (ppm)

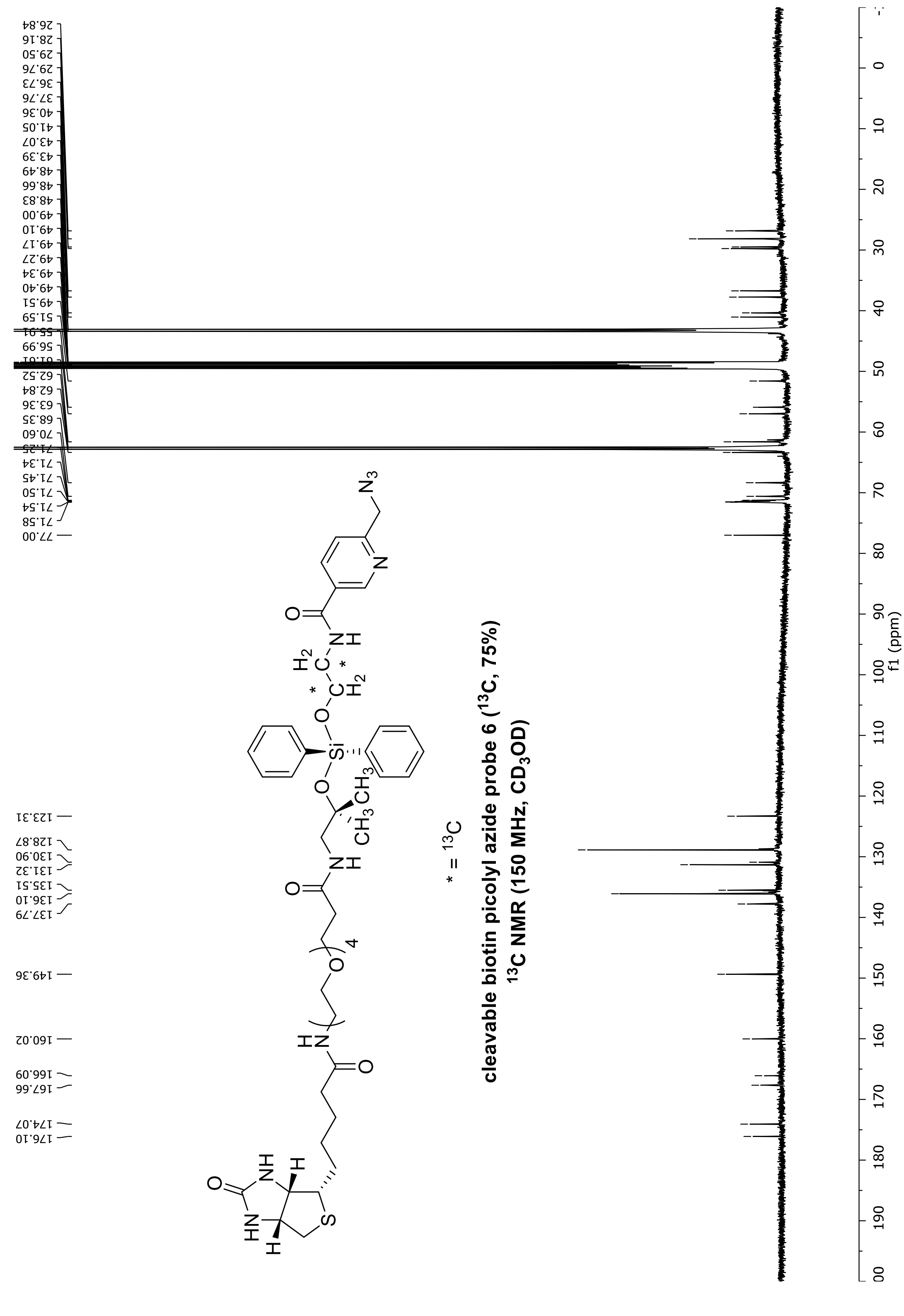
cleavable biotin picolyl azide probe 6 (13C, 75%)
13C NMR (150 MHz, CD3OD)
* = 13C
f1 (ppm)
26.84
28.16
29.50
29.76
36.73
37.76
40.36
41.05
43.07
43.39
48.49
48.66
48.83
49.00
49.10
49.17
49.27
49.34
49.40
49.51
51.59
55.91
56.99
61.01
62.52
62.84
63.36
68.35
70.60
71.25
71.34
71.45
71.50
71.54
71.58
77.00
123.31
128.87
130.90
131.32
135.51
136.10
137.79
149.36
160.02
166.09
167.66
174.07
176.10

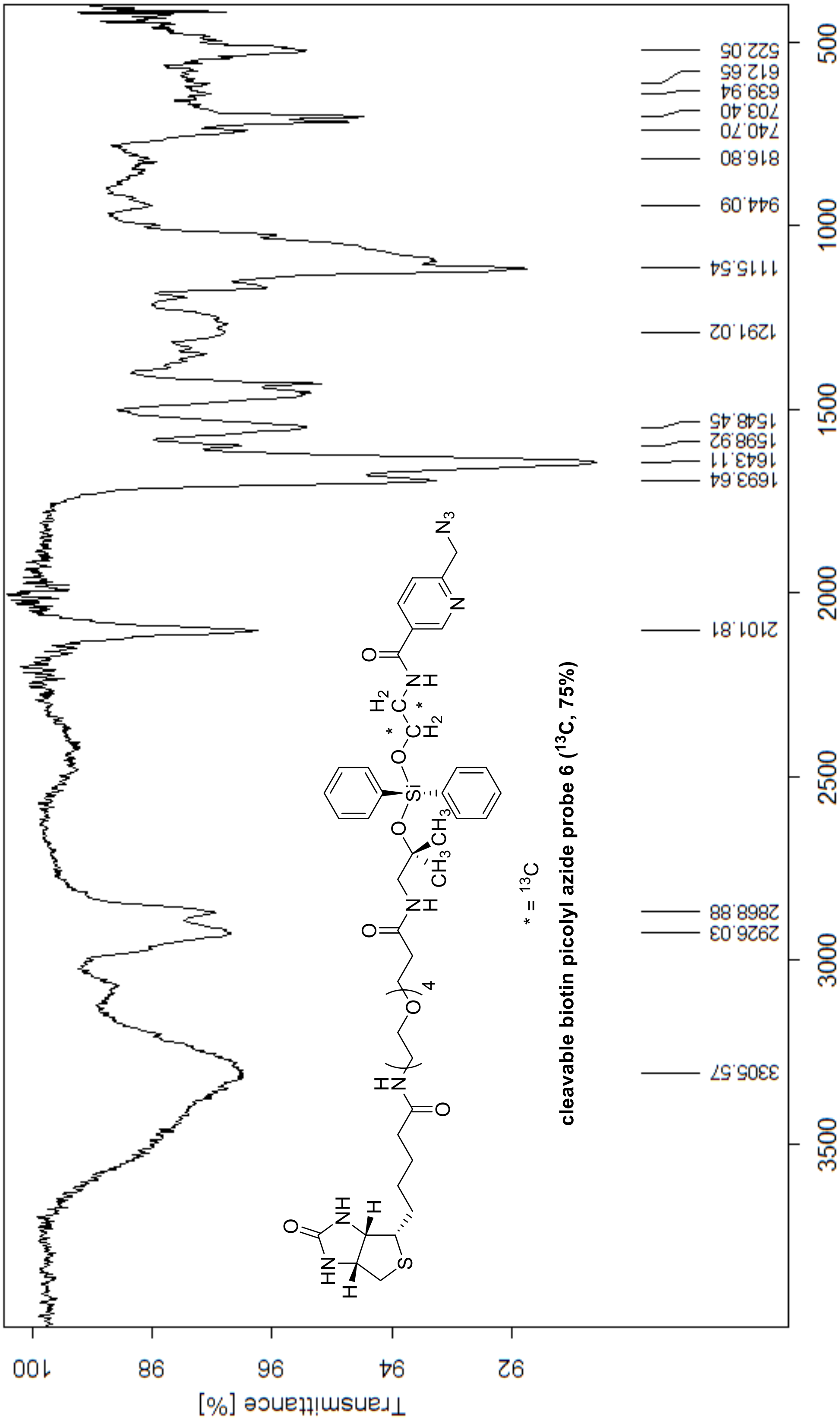
cleavable biotin picolyl azide probe 6 (13C, 75%)
* = 13C
Wavenumber cm-1
Transmittance [%]
3305.57
2926.03
2868.88
2101.81
1693.64
1643.11
1598.92
1548.45
1291.02
1115.54
944.09
816.80
740.70
703.40
639.94
612.65
522.05

**Catalog of Unique Binding Site Peptide Spectral Matches.**

Peptide sequence: HGGPQYCR
Modifications: H1(pCel); C7 (carbamidomethyl)
Protein: PTGES
Missed cleavage: 0
Charge: 2
m/z: 798.81024
MH+ [Da]: 1596.61321
RT [min]: 67.8142
First Scan: 16551
XCorr: 2.48

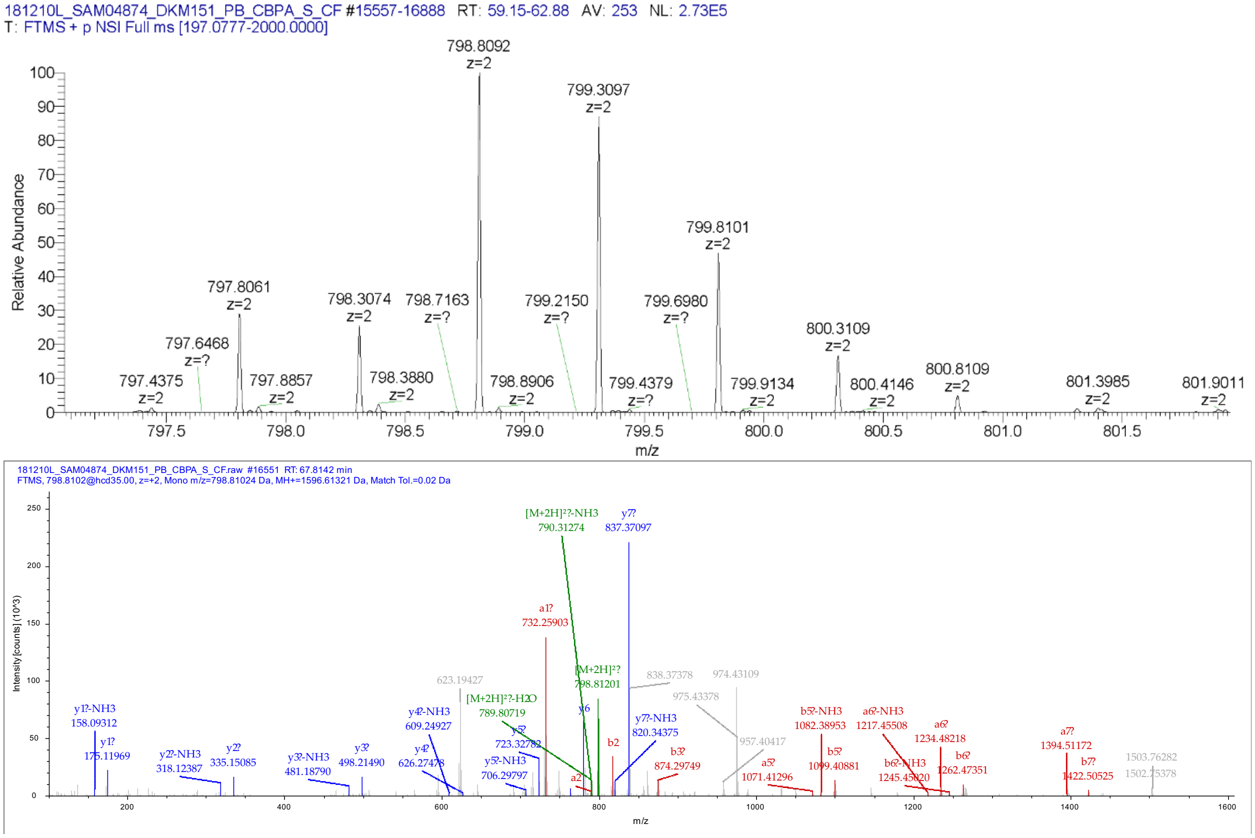

Peptide sequence: AFANPEDALRHGGPQYCR
Modifications: H11 (pCel); C17 (Carbamidomethyl)
Protein: PTGES
Missed cleavage: 1
Charge: 4
m/z: 671.04199
MH+ [Da]: 2681.14614
RT [min]: 91.0535
First Scan: 24425
XCorr: 3.54

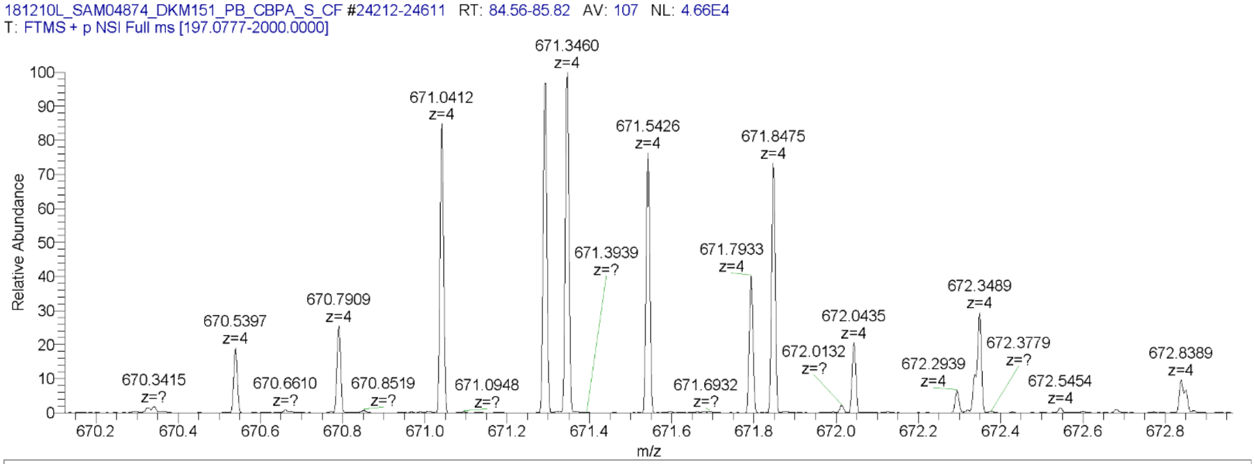


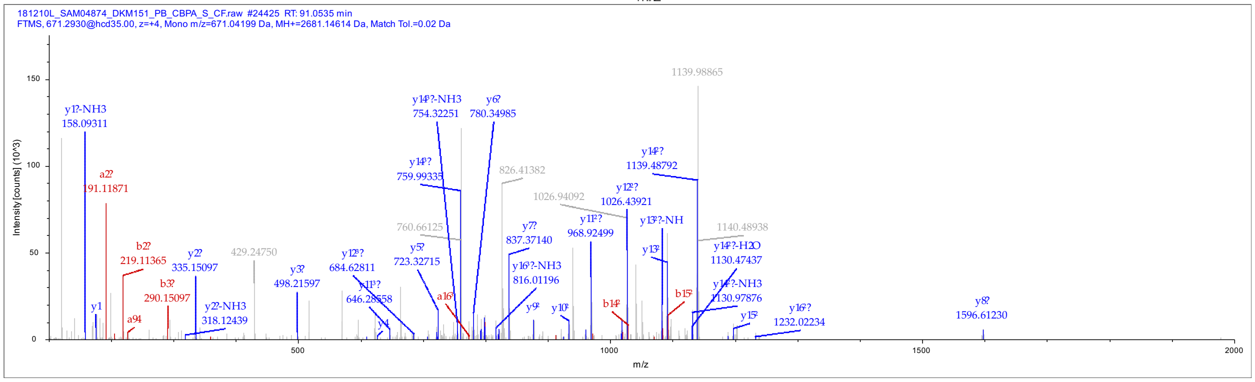

Peptide sequence: KAFANPEDALRHGGPQYCR
Modifications: H12 (pCel); C18 (Carbamidomethyl)
Protein: PTGES
Missed cleavage: 2
Charge: 4
m/z: 703.06610
MH+ [Da]: 2809.24257
RT [min]: 82.1717
First Scan: 21568
XCorr: 3.79

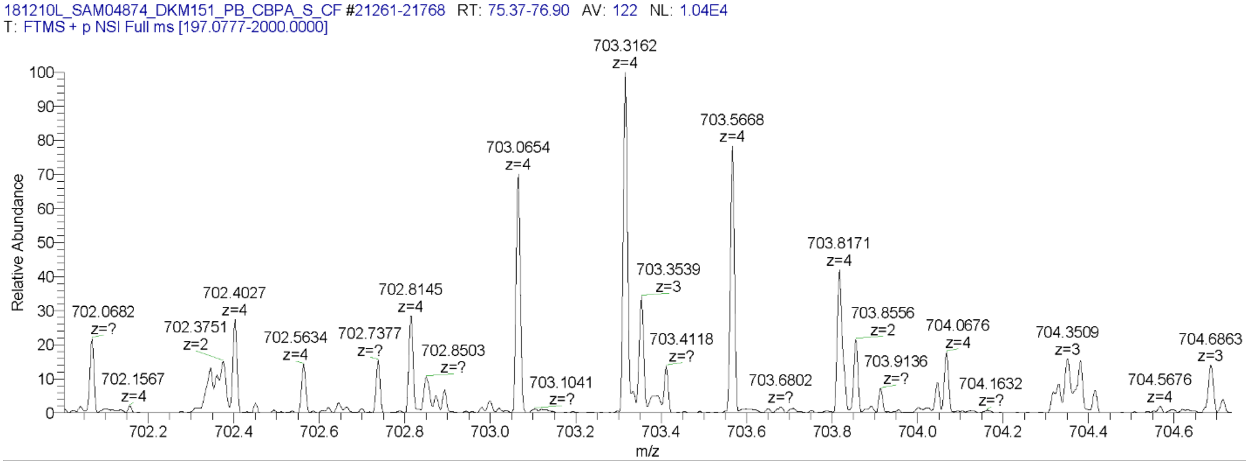


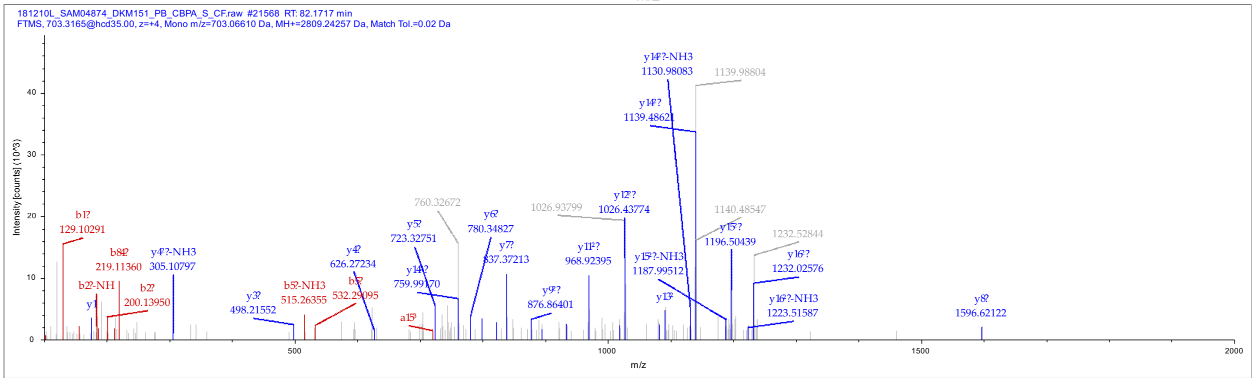

Peptide sequence: GSQGPLPFHEK
Modifications: E10 (pCel)
Protein: SLC25A10
Missed cleavage: 0
Charge: 3
m/z: 606.93524
MH+ [Da]: 1818.79117
RT [min]: 99.6125
First Scan: 25567
XCorr: 2.03

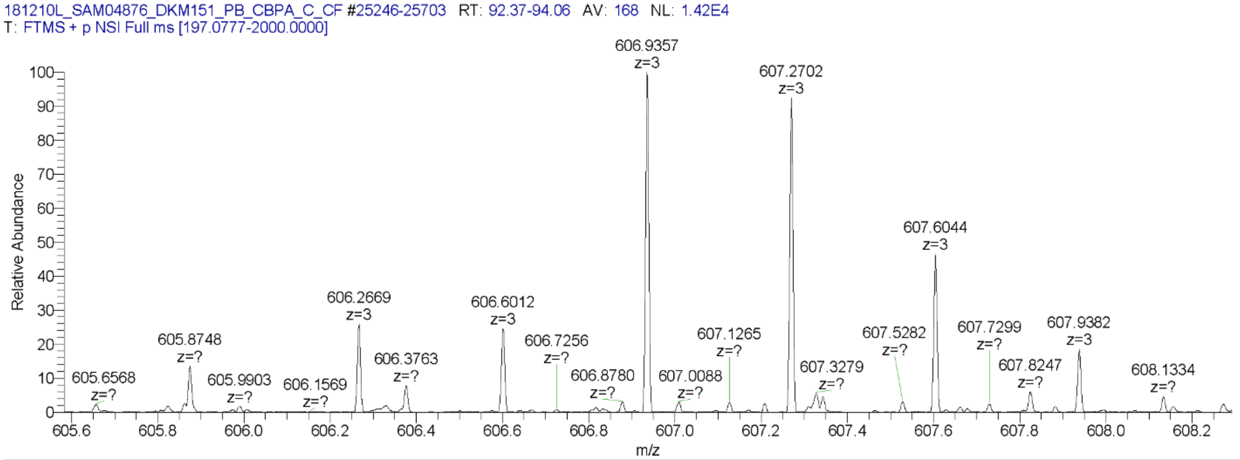


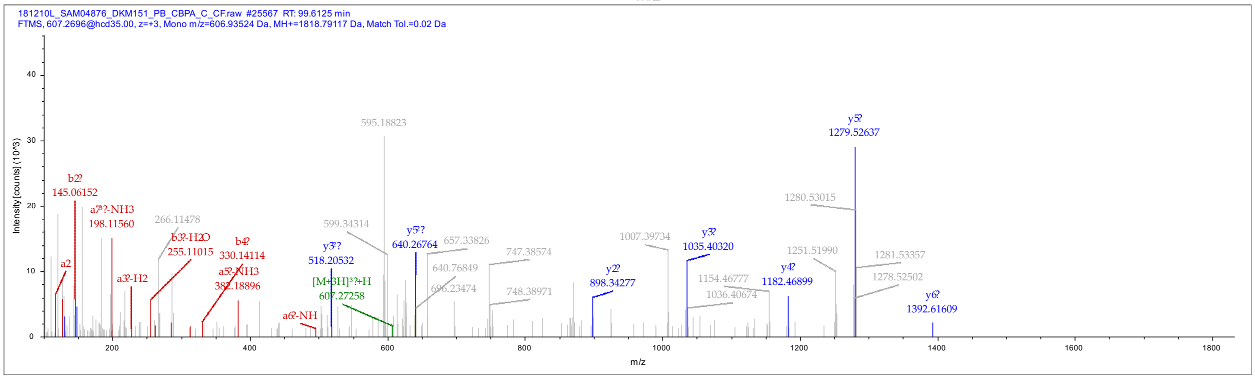

Peptide sequence: YSEGYPGKR
Modifications: S2 (pCel)
Protein: SHMT2
Missed cleavage: 1
Charge: 3
m/z: 560.23682
MH+ [Da]: 1678.69528
RT [min]: 80.2668
First Scan: 20923
XCorr: 1.59

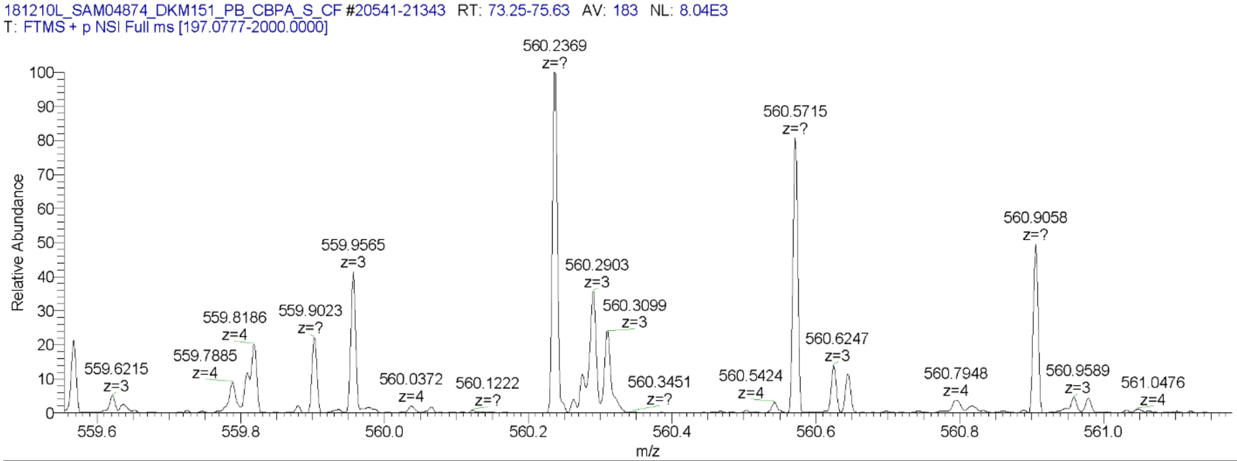


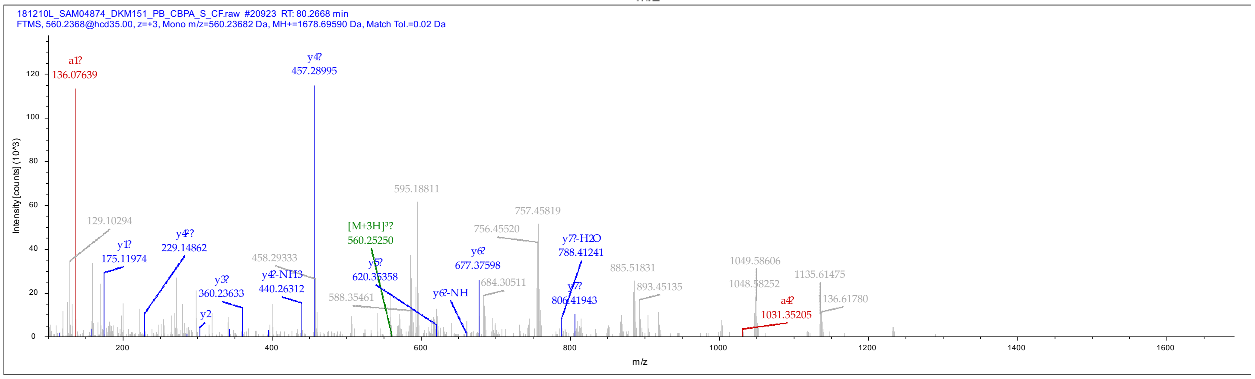

Peptide sequence: YSEGYPGK
Modifications: E3 (pCel)
Protein: SHMT2
Missed cleavage: 0
Charge: 2
m/z: 723.28613
MH+ [Da]: 1445.56499
RT [min]: 98.5954
First Scan: 20748
XCorr: 1.96

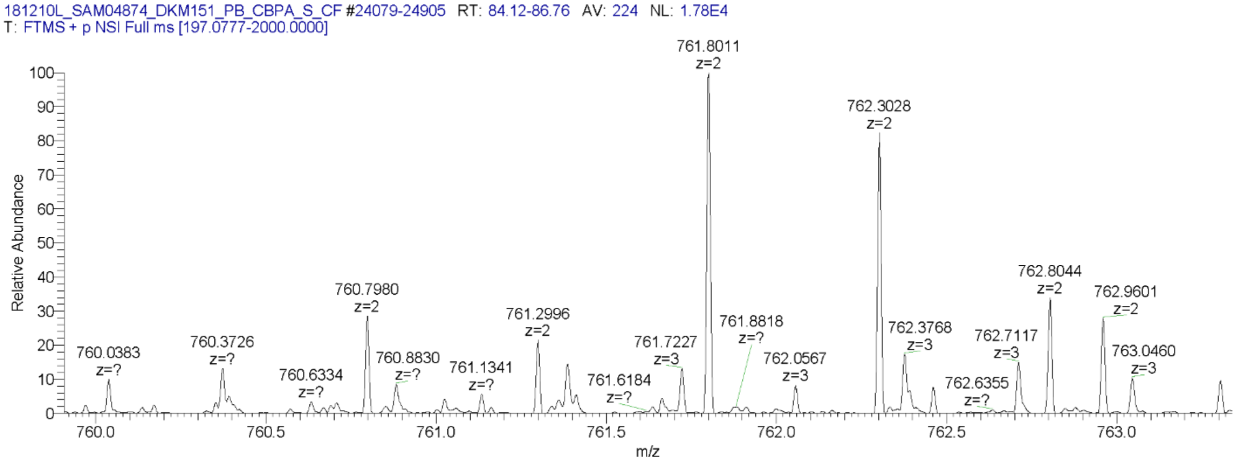


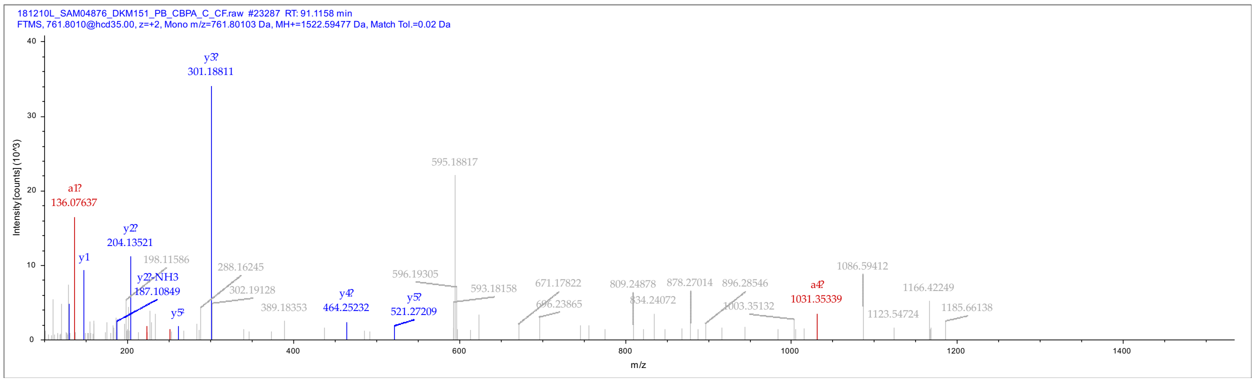

Peptide sequence: VTLELGGK
Modifications: E4 (pCel)
Protein: ALDH2
Missed cleavage: 0
Charge: 2
m/z: 719.83716
MH+ [Da]: 1438.66704
RT [min]: 106.8143
First Scan: 29319
XCorr: 1.87

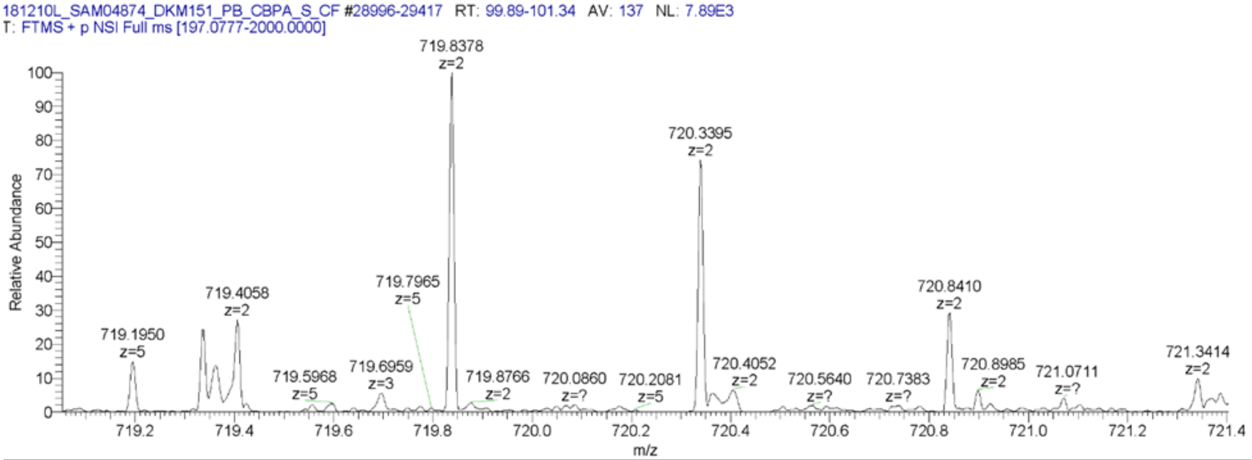


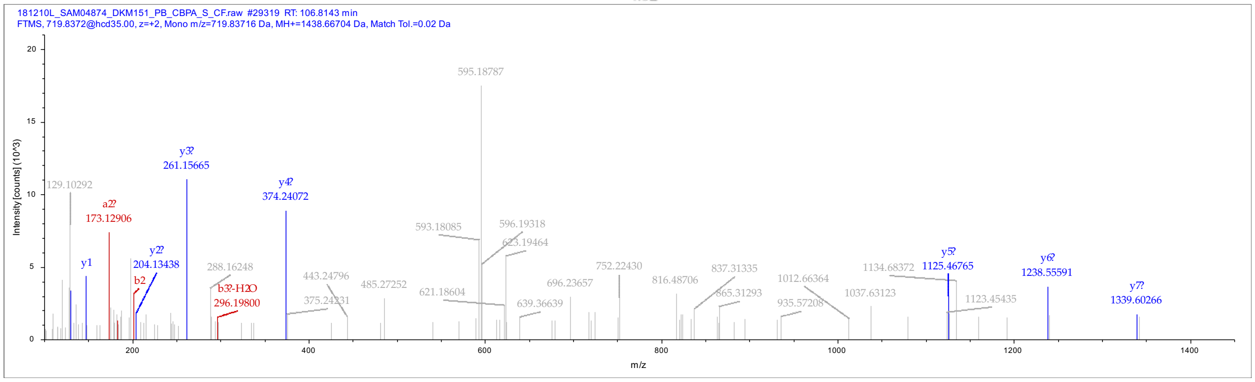

**Bibliography.**


1. Still, W. C., Kahn, M. & Mitra, A. Rapid Chromatographic Technique for Preparative Separations with Moderate Resolution. *J. Org. Chem.* **43**, 2923-2925 (1978).
2. Pangborn, A. B. *et al.* Safe and Convenient Procedure for Solvent Purification. *Organometallics* **15**, 1518-1520 (1996).
3. Lee, P. J. J. & Compton, B. J. Desctructible surfactants and uses thereof. USA patent (2007).
4. Uttamapinant, C. *et al.* Fast, Cell-compatible Click Chemistry with Copper-chelating Azides for Biomolecular Labeling. *Angew Chem Int Ed* **51**, 5852-5856 (2012).
5. Szychowski, J. *et al.* Cleavable Biotin Probes for Labeling of Biomolecules via Azide–Alkyne Cycloaddition. *J Am Chem Soc* **132**, 18351–18360 (2010).
6. Gao, J., Mfuh, A., Amako, Y. & Woo, C. M. Small Molecule Interactome Mapping by Photoaffinity Labeling Reveals Binding Site Hotspots for the NSAIDs. *J Am Chem Soc* **140**, 4259-4268 (2018).
7. Li, Z. *et al.* Design and Synthesis of Minimalist Terminal Alkyne-Containing Diazirine Photo-Crosslinkers and Their Incorporation into Kinase Inhibitors for Cell- and Tissue-Based Proteome Profiling. *Angew Chem Int Ed* **52**, 8551-8556 (2013).
8. Woo, C. M. *et al.* Development of IsoTaG, a Chemical Glycoproteomics Technique for Profiling Intact N- and O-Glycopeptides from Whole Cell Proteomes. *J Proteome Res* **16**, 1706-1718 (2017).
